# Multimodal Spatial Proteomic Profiling in Acute Myeloid Leukemia

**DOI:** 10.1101/2024.08.30.610347

**Authors:** Christopher P. Ly, Ivo Veletic, Christopher D. Pacheco, Enes Dasdemir, Fatima Z. Jelloul, Sammy Ferri-Borgongo, Akshay V. Basi, Javier A. Gomez, Jessica L. Root, Patrick K. Reville, Padmanee Sharma, Sreyashi Basu, Andres E. Quesada, Carlos Bueso-Ramos, Taghi Manshouri, Miriam Garcia, Jared K. Burks, Hussein A. Abbas

## Abstract

Acute myeloid leukemia (AML) resides in an immune rich microenvironment, yet, immune-based therapies have faltered in eliciting durable responses. Bridging this paradox requires a comprehensive understanding of leukemic interactions within the bone marrow microenvironment. We optimized a high-throughput tissue-microarray based pipeline for high-plex spatial immunofluorescence and mass cytometry imaging on a single slide, capturing immune, tumor, and structural components. Using unbiased clustering on the spatial K function, we unveiled the presence of tertiary lymphoid-like aggregates in bone marrow which we validated using spatial transcriptomics and an independent proteomics approach. We then found validated TLS signatures predictive of outcomes in AML using an integrated public 480 patient transcriptomic dataset. By harnessing high-plex spatial proteomics, we open the possibility of discovering of novel structures and interactions that underpin leukemic immune response. Further, our study’s methodologies and resources can be adapted for other bone marrow diseases where decalcification and autofluorescence present challenges.

## Introduction

Acute Myeloid Leukemia (AML) is a malignant clonal disorder of the hematopoietic system^1^. Approximately, 50% of patients achieve complete remission with induction therapy, but overall survival remains <30% at five years. Further, AML patients who relapse or are refractory to chemotherapy have dismal prognosis^2^. Immunotherapies have emerged as a promising therapeutic option in solid cancers, yet efficacy in AML has been disappointing in clinical trials^3^.

Leukemic cells have been shown to reprogram the surrounding bone marrow microenvironment (BMM) in the course of disease progression, modulating anti-tumor immune response and enabling immune evasion^4,5^. Filling the AML-immune cell interaction knowledge gap to uncover actionable therapeutics requires deeper insights on how cells within the BMM interact. The advent of high-parameter targeted spatial proteomics technologies^6–9^, which capture the spatial distribution of proteins and their expression, has allowed for a richer understanding of the tumor immune microenvironment. Non-spatial technologies such as single cell RNA sequencing and flow cytometry disaggregate cells and are unable to resolve tissue structure, discarding critical organization of cell interactions such as tertiary lymphoid structures (TLS) which, as part of the adaptive immune response, have been shown to have prognostic implications in cancer, improved response to immune checkpoint therapy^10^ and increased lymphocyte infiltration^11^.

Sequential immunofluorescence (IF) is a novel technology that allows for the high-plex interrogation of the spatial proteome. However, the technique comes with unique challenges in formalin-fixed, paraffin-embedded (FFPE) bone marrow tissue: endogenous autofluorescence, tissue and bone deformation through repeated cycles of imaging, and necessity of decalcification, which degrades morphology and immunoreactivity^12^.

Given these limitations, we developed a robust sample-to-analysis workflow to spatially capture the proteome of the BMM using an integrative combination of IF and IMC, maximizing the strengths of each modality and enabling spatial alignment at the single-cell level. Additionally, hematoxylin and eosin (H&E) staining on an adjacent slide captures structural components, granting a comprehensive view of the BMM.

We present an optimized workflow for the high-plex spatial proteomic imaging of leukemic BMM using the COMET sequential IF system to uncover the appearance of a TLS-like aggregate in AML-affected bone marrow and demonstrate the capability for discovering novel AML-BMM interactions with prognostic implications. Furthermore, we provide a resource of validated antibodies for use in bone marrow imaging. These strategies allow high throughput multimodal spatial characterization and can be applied to other bone marrow diseases beyond AML.

## Results

### An optimized high-plex spatial proteomics workflow

We developed a robust sample-to-analysis workflow optimized for speed, flexibility, and accurate recapitulation of the bone marrow (**Figure 1**). Seven bone marrow biopsies from seven adult AML patients were organized into two TMAs using an automated TMA builder for high throughput, reduced staining and imaging costs, and reduced batch effect without compromising spatial relationships^13^. Compared to manual TMA generation, the automated TMA builder provided faster turnaround time and consistent spacing, thickness, and core height. We found a 1.5 mm core diameter to be ideal for tissue adhesion as smaller cores (≤1 mm) tended to detach under our HIER protocol (see “Materials and Methods”). The resulting TMA FFPE block was sectioned, mounted onto three standard glass slides for H&E staining, proteomic imaging using the COMET IF and Hyperion Xti IMC systems, and any additional assays. The block was then archived for future use. Section thickness was optimized at 4 µm based on the depth of focus of the COMET as well as signal-to-noise ratio for IF in FFPE tissues^14^.

**Figure 1.**
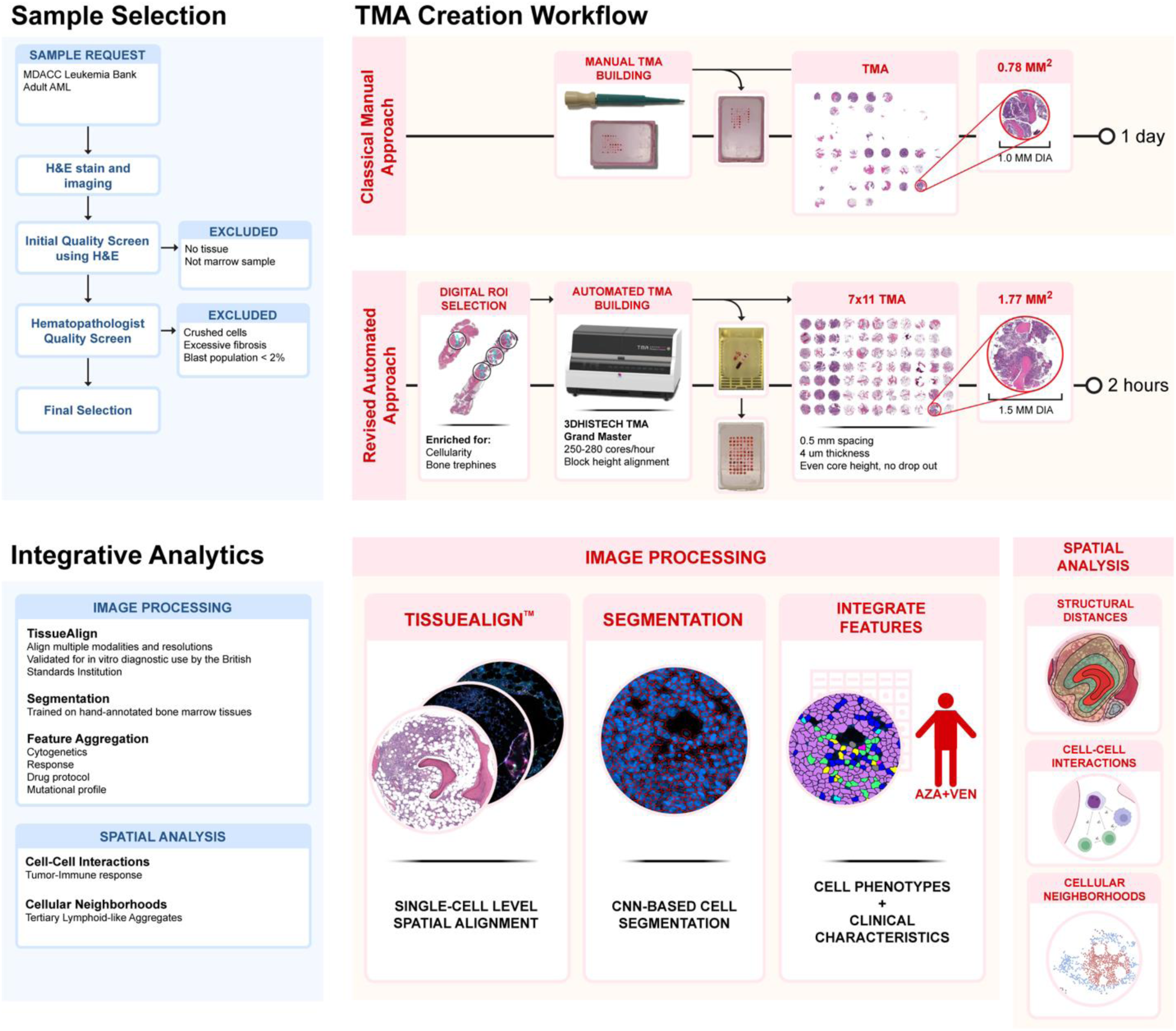
Overview of the multimodal bone marrow imaging pipeline. Patient bone marrow biopsies are ordered based on study suitability then undergo rigorous quality screening. A digital automated approach for TMA building is used, which increases quality, reproducibility, and speed compared to manual TMA building. Analysis workflow begins by aligning all modalities to a common reference. Imaging data is decomposed into cells by cell segmentation with the UNET convolutional neural network. Clinical and cell features are aggregated, allowing for comprehensive high-resolution analysis on spatial neighborhoods, structural proximity, and marker enrichment. Figure created in Biorender.

### Spatial immunofluorescence in FFPE bone marrow reliably captures the bone marrow microenvironment

Using our optimized protocol, we spatially interrogated trephine bone marrows from 7 AML patients (mean age = 67.9 years; range 62.8-72.55; 43% Male) pre-and post-AML directed therapy. This analysis involved 16 TMA cores distributed across two TMAs using the COMET approach (**Figure 2A**, characteristics detailed in **Table 1**). A total of 28 antibodies, which included canonical markers for immune cells (n = 12), AML cells (n = 4), and functional markers (n = 12), were applied to each TMA (**Table 2**). To overcome autofluorescence emerging from lipofuscin, red blood cells, and extracellular matrix components such as collagen and elastin^15,16^, we devised an algorithm to optimally subtract the autofluorescence from the imaged tissue. Briefly, we utilized a the autofluorescence quencher Trueblack and manually adjusted the magnitude of the subtraction by the average endogenous autofluorescence per cycle to obtain a scaled subtraction effect, which recovered lost signal while minimizing noise (See Supplemental Methods, **Supplemental Figure 1A-F**).

**Figure 2.**
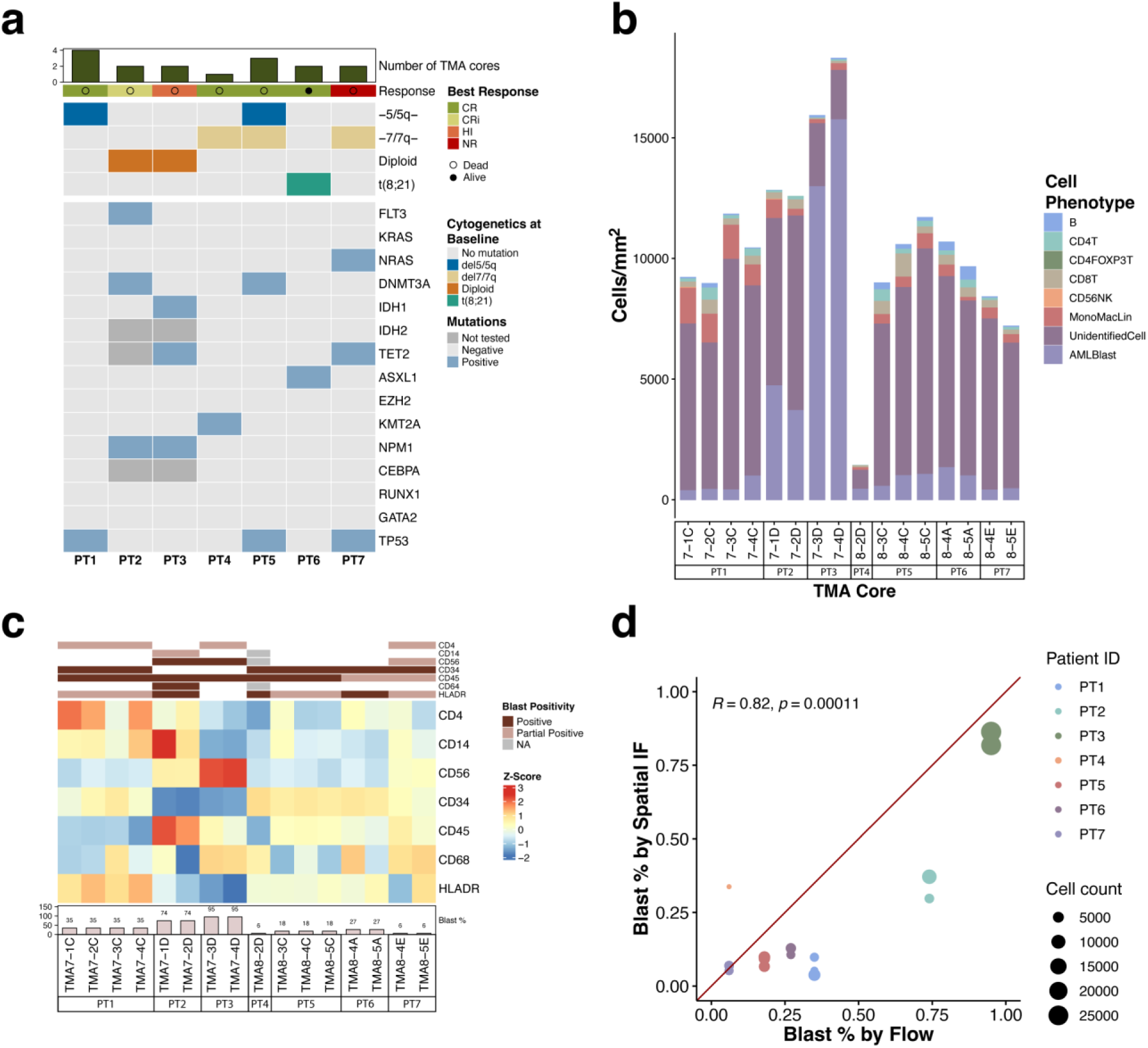
Immunofluorescent profiling of the bone marrow corresponds with clinical spectral flow. (A) Oncoprint of the cytogenetics, response, and mutational characteristics of patients 1-7. (B) Stacked bar plot showing cell type densities by TMA core. (C) Immunophenotype of AML cells by flow aligns with marker expression in spatial IF. (D) AML blasts measured by IF significantly correlate with reported blast percentages by flow cytometry (Pearson correlation, R = 0.82, p = 0.00011).

**Table 1:** Patient Characteristics. Abbreviations: AML indicates acute myeloid leukemia, MDS, myelodysplastic syndrome; RCC, renal cell carcinoma, PCa, prostate cancer; NSCLC, non-small cell lung cancer; DLBCL, diffuse large B-cell lymphoma; NHL, Non-Hodgkin’s lymphoma; BC, breast cancer; HL Hodgkin’s lymphoma; IDA, idarubicin; CLAD, cladribine; LDAC, low-dose cytarabine; VEN, Venetoclax; DEX, dexamethasone; VIN, vincristine; AZA, azacytidine; IND, investigational new drug; CR, complete remission; CRi, complete remission with incomplete count recovery; NR, no response; HI, hematologic improvement; NA, not available. *AML subtypes were determined according to the World Health Organization’s 5th edition^64^. †Mutations were detected using the 81-gene next-generation sequencing panel, EndLeukemia Assay v1.

| Characteristic (unit) | PT1 | PT2 | PT3 | PT4 | PT5 | PT6 | PT7 |
| --- | --- | --- | --- | --- | --- | --- | --- |
| Demographics |  |  |  |  |  |  |  |
| Age (years) | 72 | 69 | 68 | 63 | 66 | 62 | 72 |
| Sex/gender | Male | Female | Male | Male | Female | Female | Female |
| Race/ethnicity | White | White | White | White | Asian | Hispanic | White |
| Disease progression |  |  |  |  |  |  |  |
| AML subtype* | AML-M2 | AML-M4 | AML-M1 | AML-M1 | AML-M2 | AML-M2 | AML-M2 |
| Origin | De novo | De novo | t-AML | De novo | t-AML | t-AML | t-AML |
| Previous malignancies | None | None | RCC, PCa | None | NSCLC, MDS | DLBCL | NHL, BC, HL, MDS |
| Time since diagnosis (months) | 10 | 0 | 0 | 9 | 7 | 0 | 5 |
| Status | Deceased | Deceased | Deceased | Deceased | Deceased | Alive | Deceased |
| Survival time (months) | 17 | 9 | 2 | 11 | 11 | 9 | 9 |
| **Cytogenetic and molecular features** |  |  |  |  |  |  |  |
| Karyotype | Monosomy 5 | Diploid | Diploid | Monosomy 7 | Monosomy 5 | t(8;21) | Monosomy 7 |
| Mutations<sup>3</sup> | TP53 | NPM1, FLT3, DNMT3A | NPM1, TET2, IDH1 | KMT2A | DNMT3A, SH2B3, TP53 | ASXL1 | NRAS, TET2, TP53, SF3B1 |
| **Bone marrow counts** |  |  |  |  |  |  |  |
| Cellularity (%) | 10-20 | 95 | 100 | <5 | 30-40 | 40 | 90 |
| Lymphocytes (%) | 14 | 4 | 2 | NA | 24 | 10 | 36 |
| Blast (%) | 35 | 60 | 95 | 6 | 18 | 27 | 6 |
| **Peripheral blood counts** |  |  |  |  |  |  |  |
| WBC (×10<sup>6</sup>/L) | NA | 41.9 | 4.98 | NA | 0.86 | 0.17 | 0.99 |
| Lymphocytes (%) | NA | 19 | 10 | NA | 54 | 61 | 78 |
| Monocytes (%) | NA | 50 | NA | NA | 6 | 5 | 5 |
| **Treatment** |  |  |  |  |  |  |  |
| Active agents | SGI 110+IDA | Treatment naïve | Treatment naïve | CLAD+LDAC+VEN+DEX+VIN | Hu5F-G4+AZA | Treatment naïve | AZA+IND |
| Pre/Post Treatment | Post | Pre | Pre | Post | Post | Pre | Post |
| Response category | CR | CRi | NR | CR | CR | CR | HI |

**Table 2:** COMET Antibodies.

| COMET Antibodies |  |  |  |  |  |  |  |  |
| --- | --- | --- | --- | --- | --- | --- | --- | --- |
| Cycle | Channel | Antigen | Clone | Clonality | Source | Catalog ID | Dilution | Quality |
| 1 | Cy5 | CD34 | EP373Y | Rabbit Monoclonal | Abcam | ab81289 | 1:50 | *** |
|  | TRITC | CD68 | KP1 | Mouse Monoclonal | Bethyl | A500-018A | 1:100 | *** |
| 2 | Cy5 | CX3CR1 | POLY | Rabbit Polyclonal | Invitrogen | 14-6093-81 | 1:50 | *** |
|  | TRITC | GRZK | 1C3A4 | Mouse Monoclonal | ProteinTech | 67272-1-Ig | 1:100 | * |
| 3 | Cy5 | BCL11a | BLR073G | Rabbit Monoclonal | Bethyl | A700-073 | 1:50 | *** |
|  | TRITC | CD3 | 3F3A1 | Mouse Monoclonal | ProteinTech | 60181-1-Ig | 1:100 | * |
| 4 | Cy5 | HLADRPQ | CR3-43 | Mouse Monoclonal | Bethyl | A500-034A | 1:50 | ** |
|  | TRITC | CD11c | D3V1E | Rabbit Monoclonal | Cell Signaling Technology | 93233 | 1:100 | ** |
| 5 | Cy5 | IFITM3 | POLY | Rabbit Polyclonal | ProteinTech | 11714-1-AP | 1:50 | *** |
|  | TRITC | CD74 | PIN.1 | Mouse Monoclonal | NovusBio | NBP2-80657 | 1:200 | ** |
| 6 | Cy5 | CD4 | BL-155-1C11 | Rabbit Monoclonal | Bethyl | A700-015 | 1:50 | *** |
|  | TRITC | CD20 | L26 | Mouse Monoclonal | Bethyl | A500-017A | 1:100 | *** |
| 7 | Cy5 | CD8a | C8/144B | Mouse Monoclonal | Bethyl | A500-021A | 1:100 | ** |
|  | TRITC | GRZB | POLY | Rabbit Polyclonal | ProteinTech | 13588-1-AP | 1:250 | ** |
| 8 | Cy5 | BST2 | 26F8 | Mouse Monoclonal | Invitrogen | 16-3179-82 | 1:50 | * |
|  | TRITC | FOXP3 | BLR034F | Rabbit Monoclonal | Bethyl | A700-034 | 1:100 | ** |
| 9 | Cy5 | CD14 | BLR187J | Rabbit Monoclonal | Bethyl | A700-187 | 1:50 | ** |
|  | TRITC | HLAE | 1A4G3 | Mouse Monoclonal | ProteinTech | 66530-1-Ig | 1:200 | *** |
| 10 | Cy5 | CD33 | PWS55 | Mouse Monoclonal | CellMarque | 133M14 | 1:25 | * |
|  | TRITC | CD45 | BL-178-12C7 | Rabbit Monoclonal | Bethyl | BL-178-12C7 | 1:800 | ** |
| 11 | Cy5 | CD34 | POLY | Rabbit Polyclonal | ProteinTech | 14486-1-AP | 1:100 | ** |
|  | TRITC | P53 | 6C4B6 | Mouse Monoclonal | ProteinTech | 60283-2-Ig | 1:100 | * |
| 12 | Cy5 | Ki67 | EPR3610 | Rabbit Monoclonal | Abcam | ab92742 | 1:100 | *** |
|  | TRITC | NF-kB | L8F6 | Mouse Monoclonal | Cell Signaling Technology | 6956 | 1:250 | * |
| 13 | Cy5 | CD56 | BLR171J | Rabbit Monoclonal | Bethyl | A700-152 | 1:500 | *** |
|  | TRITC | VEGFR2 | IG2A8 | Mouse Monoclonal | ProteinTech | 67407-1-Ig | 1:100 | * |
| 14 | Cy5 | CD163 | 10D6 | Mouse Monoclonal | Invitrogen | MA5-11458 | 1:50 | *** |
|  | TRITC | FOXP3 | D2W8E | Rabbit Monoclonal | Cell Signaling Technology | 98377 | 1:50 | * |

Out of the segmented 116,488 cells, 63,738 (54.7%) cells had at least one marker that allowed classification/identification (**Table 5**). The remaining cells likely constituted stromal cells, mesenchymal cells, or other immune subsets (such as plasma cells).

Protein expression for each cell type is shown in **Supplemental Figure 2A**. Cell types were visualized using dimensionality reduction techniques to identify each cell type (**Supplemental Figure 2B, C**). Cell densities for each cell type remained consistent within each patient between cores (**Figure 2B**). AML immunophenotype was also consistent with expression by flow cytometry (**Figure 2C**). Blast percentage significantly correlated (r=0.82; p=0.00011) with flow cytometry done as part of the clinical diagnostic work-up (**Figure 2D**). Overall, spatial IF was able to reliably capture cell populations within the bone marrow microenvironment of AML patients.

### Non-destructive sequential IF allows for proteomic validation using the same slide

To further explore other modalities to guide in identification of the AML immune microenvironment, we applied spatial imaging mass cytometry (IMC) which uses metal-labeled tagged proteins to avoid spectral overlap and image multiple analytes simultaneously. We applied a panel of 25 markers (see **Table 4**) on the same slide that we previously performed IF imaging, taking advantage of the non-destructive nature of the COMET to expand the panel and validate the findings of the COMET.

**Table 3:** Opal Antibodies.

| Opal IF |  |  |  |  |  |  |  |  |
| --- | --- | --- | --- | --- | --- | --- | --- | --- |
| Opal Fluorophore | Catalog ID | Antigen | Clone | Clonality | Source | Catalog ID | Fluorophore Dilution | Ab Dilution |
| 520 | FP1487701KT | CD4 | BL-155-1C11 | Rabbit | Bethyl | A700-15 | 1:400 | 1:250 |
| 480 | FP150001KT | CD20 | L26 | Mouse | CellMarque | 120M-85 | 1:800 | 1:250 |
| 570 | FP1488001KT | CD3 | SP7 | Rabbit | Abcam | ab16669 | 1:200 | 1:250 |
| 620 | FP1495001KT | CD8 | BLR044F | Rabbit | Bethyl | A700-044 | 1:400 | 1:250 |
| 690 | FP1497001KT | CD34 | EP373Y | Rabbit | Abcam | ab81289 | 1:200 | 1:2500 |
| DIG/780 | FP1501001KT | HLA-E | 1A4G3 | Mouse | ProteinTech | 66530-1-Ig | 1:100/1:25 | 1:200 |

**Table 4:** IMC Antibodies.

| IMC |  |  |  |  |  |  |  |
| --- | --- | --- | --- | --- | --- | --- | --- |
| Mass Tag | Antigen | Clone | Clonality | Source | Catalog ID | Dilution | Quality |
| 141Pr | CD34 | QBEND/10 | Mouse Monoclonal | Invitrogen | MA1-10202 | 1:400 | *** |
| 142Nd | T-bet | 4B10 | Mouse Monoclonal | Abcam | ab91109 | 1:100 | * |
| 144Nd | CD14 | EPR3653 | Rabbit Monoclonal | Std BioTools | 3144025D | 1:100 | ** |
| 145Nd | T-bet | D6N8B | Rabbit Monoclonal | Std BioTools | 3145015D | 1:100 | ** |
| 146Nd | CD16 | DJ130c | Mouse Monoclonal | Invitrogen | MA1-84008 | 1:100 | * |
| 147Sm | gH2AX | BLR053F | Rabbit Monoclonal | Bethyl | A700-053CF | 1:100 | ** |
| 148Nd | CD163 | BLR087G | Rabbit Monoclonal | Bethyl | A700-087CF | 1:200 | *** |
| 151Eu | CD31 | EPR3094 | Rabbit Monoclonal | Std BioTools | 3151025D | 1:200 | ** |
| 152Sm | CD45 | MEM-28 | Mouse Monoclonal | Abcam | ab8216 | 1:100 | * |
| 153Eu | CD44 | BLR038F | Rabbit Monoclonal | Bethyl | A303-872A | 1:200 | * |
| 155Gd | IDO1 | BLR031F | Rabbit Monoclonal | Bethyl | A700-031 | 1:100 | * |
| 156Gd | CD4 | EPR6855 | Rabbit Monoclonal | Std BioTools | 3156033D | 1:100 | *** |
| 159Tb | CD68 | KP1 | Mouse Monoclonal | Std BioTools | 3159035D | 1:400 | *** |
| 160Gd | GRZK | 1C3A4 | Mouse Monoclonal | ProteinTech | 67272-1-Ig | 1:100 | *** |
| 161Dy | IFITM3 | Poly | Rabbit Polyclonal | ProteinTech | 11714-1-AP | 1:50 | ** |
| 162Dy | CD8 | D8A8Y | Rabbit Monoclonal | Std BioTools | 3162035D | 1:200 | *** |
| 163Dy | BCL11a | EPR14943-44 | Rabbit Monoclonal | Abcam | ab191401 | 1:50 | *** |
| 165Ho | CD117 | Poly | Rabbit Polyclonal | ProteinTech | 18696-1-AP | 1:100 | * |
| 166Er | CD74 | LN2 | Mouse Monoclonal | Std BioTools | 3166025D | 1:100 | ** |
| 167Er | GZRB | EPR20129 | Rabbit Polyclonal | Std BioTools | 3167021D | 1:200 | ** |
| 168Er | Ki67 | SP6 | Rabbit Monoclonal | CellMarque | 275R-14 | 1:200 | *** |
| 169Tm | HLADRPQ | CR3/43 | Mouse Monoclonal | Bethyl | A500-034A | 1:100 | ** |
| 170Er | CD3 | MRQ-39 | Rabbit Monoclonal | CellMarque | 103R-94 | 1:100 | *** |
| 171Yb | CD27 | BLR083G | Rabbit Monoclonal | Bethyl | A700-083 | 1:100 | ** |
| 174Yb | HLA-E | 1A4G3 | Mouse Monoclonal | ProteinTech | 66530-1-Ig | 1:400 | *** |

We first tested whether imaging the tissue with sequential IF would degrade IMC staining. We selected two of the previously COMET-imaged TMA cores (TMA7-3C and TMA7-4C) and compared the staining when imaged subsequently imaged on the same slide using IMC. Antibody staining was highly comparable, and markers targeting the same antigen colocalized well, showcasing retained antigenicity and robustness of alignment (**Figure 3A**). Correlation analysis showed markers with high signal to background such as CD163, CD34, and Ki67 tended to cluster closely together with their imaging counterpart, as well markers with expected colocalization such as CD3, CD8, and CD45 (**Figure 3B**). Nuclear staining showed some degradation between modalities and affected cell segmentation, leading us to use the DAPI staining for nuclear segmentation going forward (**Supplemental Figure 3A-B**).

**Figure 3:**
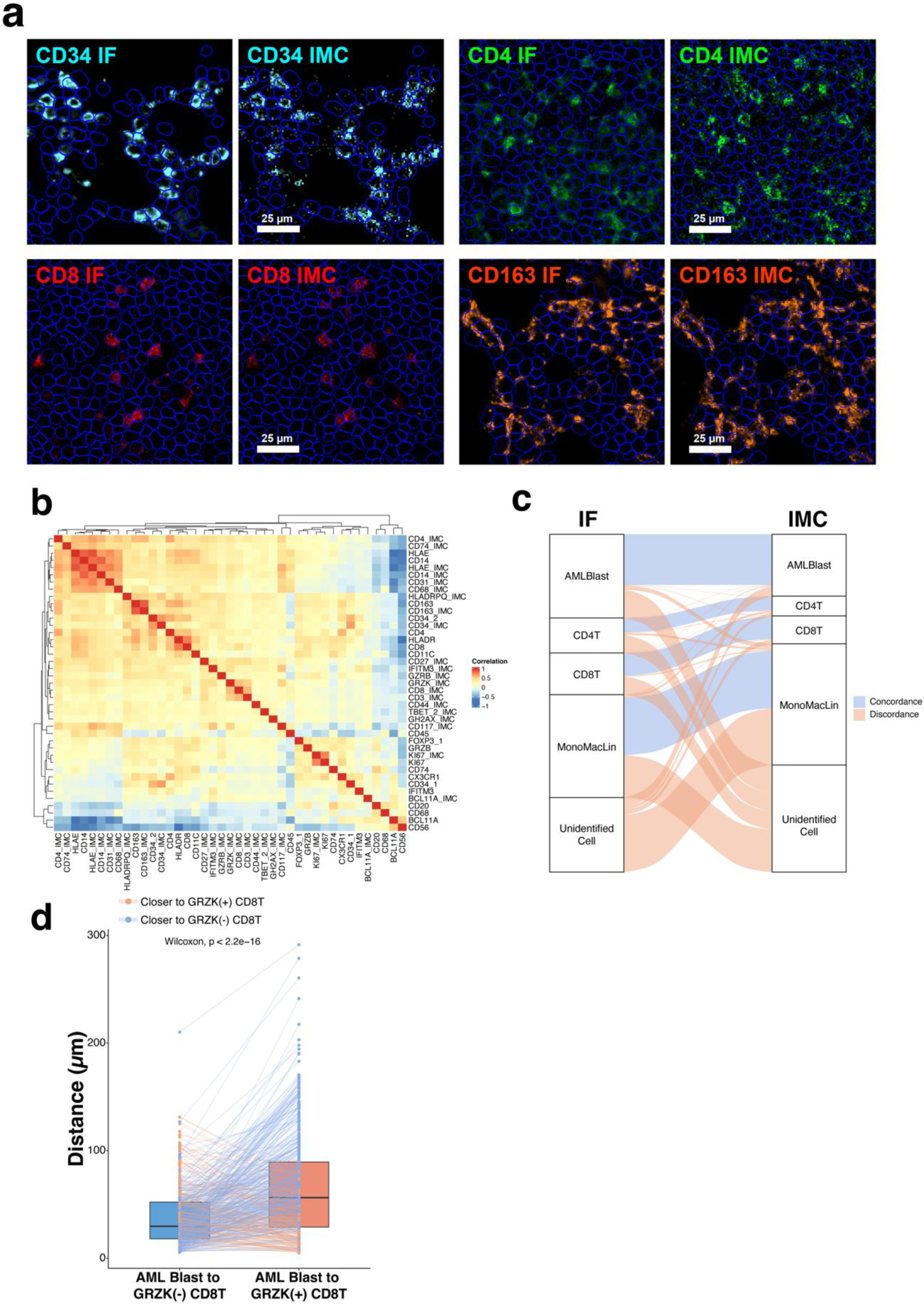
Spatial IF to IMC results in degradation of nuclear signal in IMC without compromising antigen signal: (A) Comparison of CD34 (cyan), CD4 (green), CD8 (red), and CD163 (orange) performance in IF and IMC shows high concordance, validating cell segmentation and spatial alignment approaches. (B) Correlation of same slide imaging between IF and IMC markers. (C) Alluvial plot showing proportions of concordance and discordance of the same cells phenotyped using only IMC markers and using only IF markers. (D) Distance of AML cells to granzyme K negative and granzyme K positive cells. Connected dots represent the distances for the same cell. (Paired Wilcoxon test, p < 2.2e-16).

To further evaluate IF to IMC performance, we separately phenotyped the cells iteratively captured from IF and IMC to compare how cell assignment differed. Canonical markers retained expected expression across cell types in both modalities (**Supplemental Figure 3A**). Identities of CD34(+) blasts, and CD8(+) and CD4(+) T cells were generally concordant, with IMC displaying higher specificity (82%, 79%, and 71% concordance of IMC positive populations with IF, respectively) than IF (60%, 52%, and 40% concordance) (**Figure 3C**). Monocyte/macrophage markers were more discordant, with less than 50% of cells retaining phenotypic assignment between modalities (IMC 47% concordance, IF 41% concordance). Since macrophages have large, irregular membranes that often overlap with other cells, the phenotyping is more sensitive to differences in intensity thresholds, cell segmentation, and alignment. This issue is exacerbated in nucleus-based cell segmentation, which captures morphology of small, round cells like lymphocytes better than large, irregular, or multinucleated cells. Overall, results on IMC were able to validate our findings in IF, and we found concordance between the two modalities to high enough to allow for possible panel expansion using this method in the future.

With subsequent IMC staining, we were able to image additional markers without increasing COMET cycle count. We previously published that a subset of stem-like/memory CD8T cells expressing granzyme K (GZMK) was enriched in responders to immunotherapy in AML^17^. We hypothesized AML cells could have a spatial relationship with GZMK(+) CD8(+) T cells. Using the expanded panel provided by IMC, we found that CD34(+) AML cells were significantly closer to GZMK(-) CD8(+) T cells than GZMK(+) CD8T(+) cells (**Figure 3D**). This could possibly indicate an immune-evasive response to T cells with memory-like properties, which are more likely to trigger bystander activation in inflammatory environments^18^.

### Spatial region analysis unveils tertiary lymphoid-like aggregates in adult AML

Recent developments in spatial neighborhood analysis have allowed for the identification of regions based on patterns of local cell proximity, identifying novel areas of interaction and structure with potential prognostic/therapeutic implications^19^. To test this in AML, we applied unbiased clustering over Ripley’s K function, a statistical measure of the spatial dispersion of each cell type modeled as a local point process. This revealed five distinct regions distinguished by different proportions of cell types, suggesting unique cell-cell interactions (**Figure 4A, B**). Region_1 was highly enriched in AML blasts and significantly correlated to flow cytometry blast percentage (R=0.92, p=3e-7) (**Supplemental Figure 4B).** Region_2 was primarily enriched in lymphocytes and cells of monocyte and macrophage lineage. Region_3 had a high concentration of B cells, CD4(+) T cells, CD8(+) T cells, and NK cells. Region_4 was an even mix of cell types, and Region_5 captured mainly unidentified cells. Region_4 was largest in area, as expected from the unstructured liquid formation of the BMM. Interestingly, Region_3 only appeared in tumor cores of 3/7 patients (TMA7-3C, TMA8-3C, and TMA8-5E) and consisted of a tight cluster of B and T lymphocytes. These aggregates varied in size and organization, with TMA7-3C having the largest aggregate and rough organization of a B/CD4 T cell core and CD4/CD8 T cell border, while TMA8-3C and TMA8-5E were much smaller and contained a thoroughly mixed B and T cell population. This finding was consistent of various states of maturity of TLS found in solid tumors based on composition and organization (**Figure 4C, D**). B cells in region_3 also displayed significantly higher expression of Ki67 compared to other regions, possibly indicative of a proliferative B cell core, a previously reported characteristic of TLS in solid cancers (**Supplemental Figure 4C**). We then stratified patients by presence of region_3 aggregates. Looking at differences between cell densities, we were surprised to find that patients with region_3 aggregates had significantly higher densities monocyte and macrophage lineage cells, and while T and B cells were overall higher in density, they were not significantly so (**Figure 5A**). Patients with region_3 aggregates also exhibited a significantly higher percentage of CD4T and CDT cells in the AML-enriched region_1, analogous increased tumor lymphocyte infiltration in solid tumors (**Figure 5B**). We then were able to validate the presence of these aggregates through the 7-color (see **Table 4**) Opal multiplex IF assay by staining subsequent sections of the original TMA blocks (**Figure 5C, Supplemental Figure 4C**). TLS presence in AML remains poorly studied, with the majority of research focusing on solid tumors. However, recently a study has described these clusters of lymphocytes as “TLS-like aggregates” in a pediatric AML cohort^20^. As such, we chose to adapt this terminology to reflect the lack of conclusive definitions of TLS in AML.

**Figure 4:**
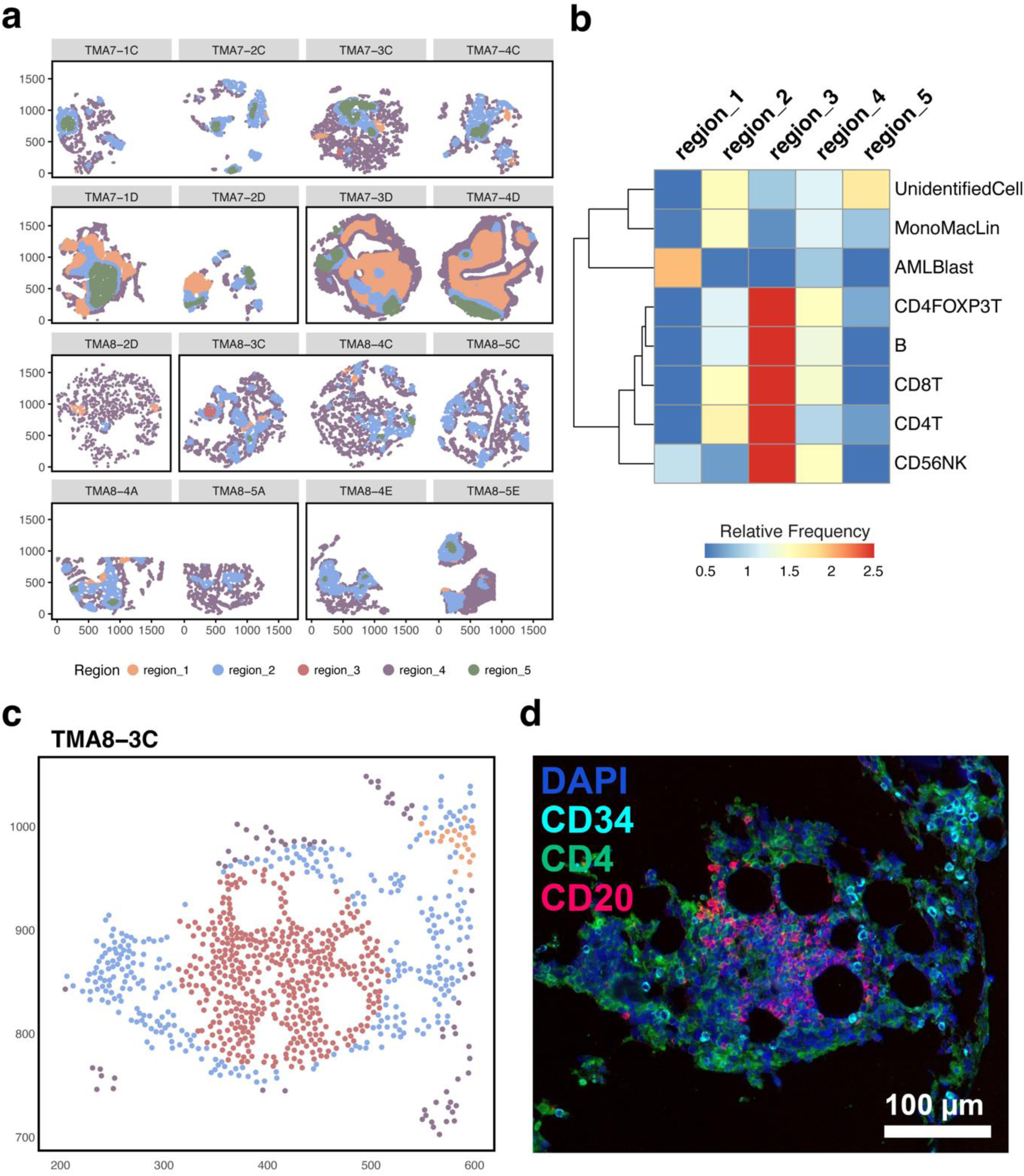
Spatial neighborhoods reveal tertiary lymphoid-like structures. (A) Unbiased clustering of the spatial K function for each cell type reveals 5 different cellular region phenotypes. (B) Relative enrichment of cell types for each cellular region. (C) Region assignment of cells in the TLS-like structure. (D) Micrograph of TLS-like structure. Blue = DAPI, cyan = CD34, green = CD4, red = CD20. Scale bar 100 µm.

**Figure 5:**
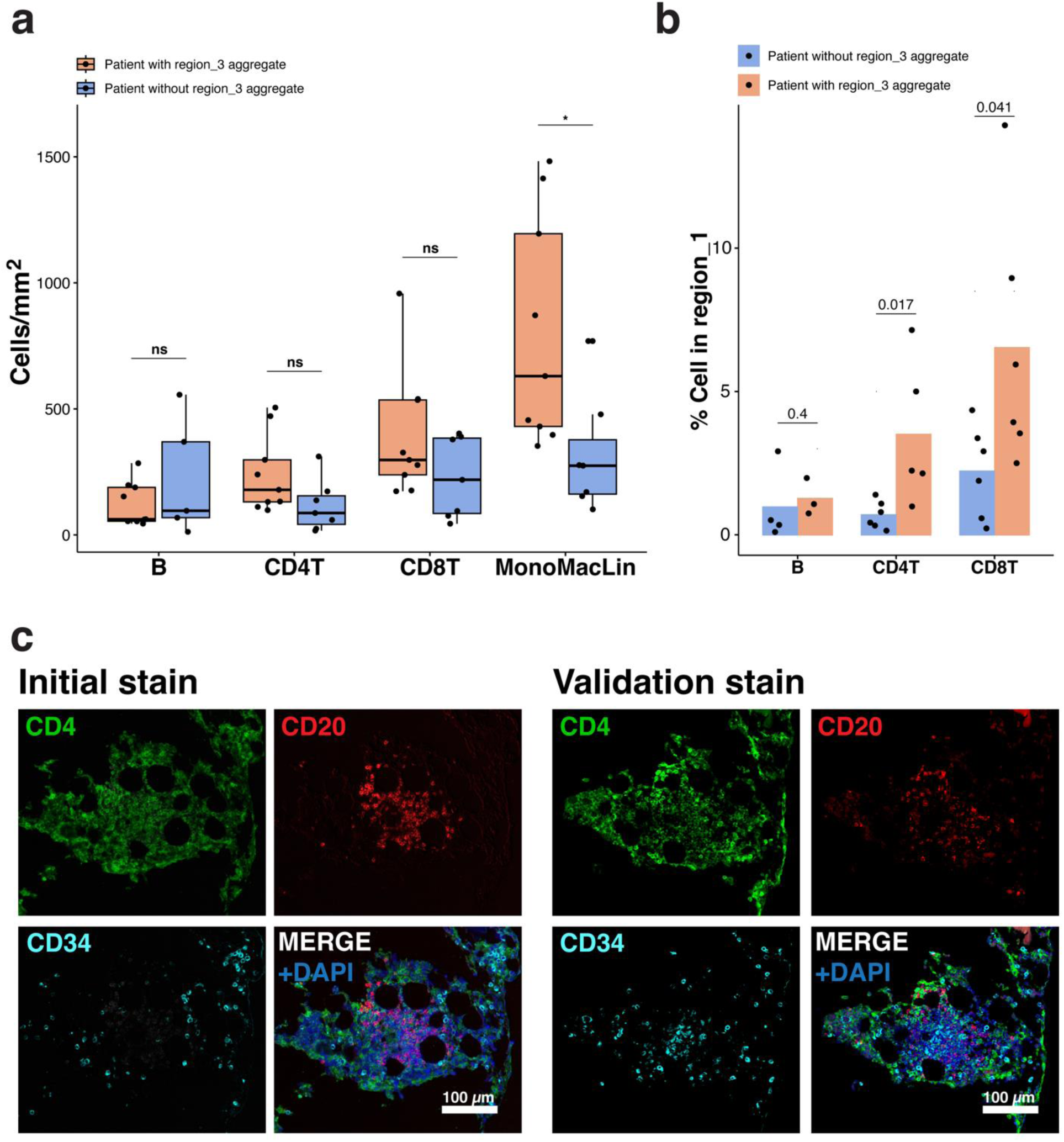
Spatial analysis reveals distance dependent AML blast responses to immune cells. (A) Cell densities for B, CD4T, CD8T, and Monocyte Macrophage lineage cells. Each dot represents an individual TMA core (Wilcoxon test, unpaired, two-sided).* = p < 0.05, ** = p < 0.01, ns = not significant). (B) Percent of B cells, CD8 T cells, and CD4 T cells in region 1 (AML enriched) areas between TMA cores with and without region_3 aggregates. Dots represent an individual TMA core. (Wilcoxon test, unpaired, two-sided). (C) COMET stain and Opal Validation stain for the region_3 aggregate of TMA8-3C.

In summary, by clustering local measures of spatial association, we were able to spatially identify a cluster of cells, denoted by region_3, of lymphocyte aggregates that resemble various stages of TLS formation and maturity in AML. These aggregates were further validated by the Opal assay.

### TLS signatures identified in Bulk RNA-seq and Spatial transcriptomics AML datasets are correlated with prognosis

Though we were able to discover TLS-like aggregates with the COMET, spatial proteomics is relatively low plex compared to RNA-seq, and our panel excluded many functional TLS markers. Using 6 published gene signatures for TLS^21–26^ (**Supplemental Table X**), we investigated the presence of TLS in AML using the Visium (10X Genomics, Pleasanton, CA) spatial transcriptomics platform, which can capture the transcriptome of the AML BMM a resolution of 55 µm spots. We focused on the signature from Cabrita et al. as it is derived from a compendium of TLS genes^21^ and not cancer specific. We performed the Visium assay on an archived bone marrow sample with a positive lymphocyte aggregate by H&E stain which overlapped with localized enrichment of the TLS signature by Cabrita et al. as well as deconvolution signatures of T cells and B cells, further suggesting TLS-like properties for these lymphocyte aggregates in AML (**Figure 6A**).

**Figure 6:**
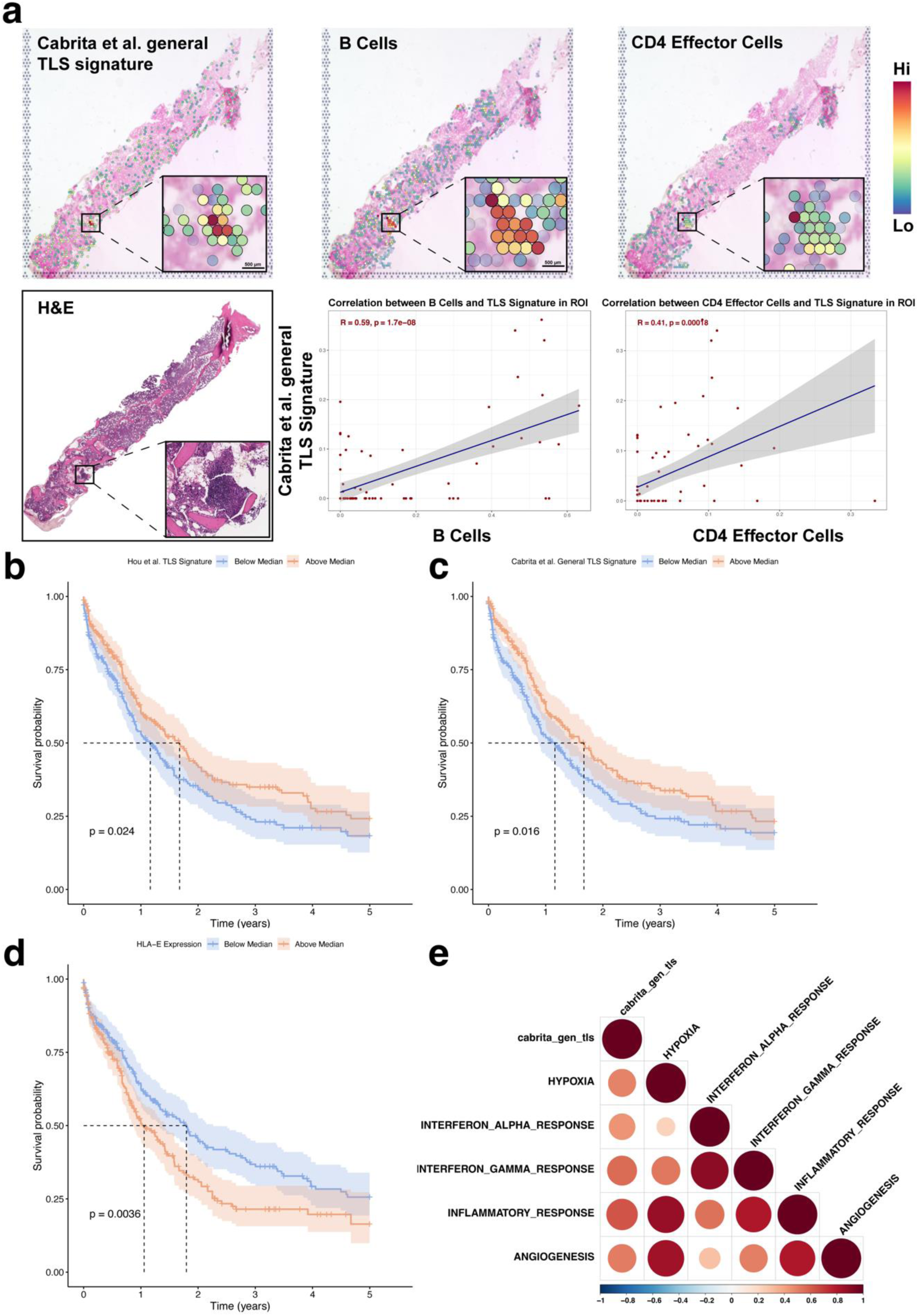
TLS signatures show prognostic value in transcriptomics. (A) Spatial transcriptomics of Cabrita et al. TLS signature, a B cell score, and CD4 T effector score in the same ROI and their correlations (Pearson correlation test). (B) Kaplan-Meier survival curves of AML patients from TCGA, Beat-AML, and MDACC by median score of the Hou et al. TLS signature. (C) Kaplan-Meier survival curves of AML patients from TCGA, Beat-AML, and MDACC by median score of the Cabrita et al. TLS signature. (D) Kaplan-Meier survival curves of AML patients from TCGA, Beat-AML, and MDACC by median HLA-E expression. (E) Correlation plot of Cabrita et al. TLS signature and significantly correlated Hallmark signatures (Pearson correlation test).

We further investigated the presence of TLS in AML using a bulk RNA-seq dataset integrating data from TCGA, MDACC, and BEAT-AML1/2 datasets and found signatures published by Cabrita et al., Hou et al. to be predictive of overall survival (**Figure 6B-C**). Additionally, in the 3/4 of the datasets, all 6 signatures significantly correlated with HLA-E expression, which we previously identified as a marker for immune evasion in AML^27^, and was significantly indicative of survival in the combined bulk RNA dataset (**Supplemental Figure 5A, Figure 6D**). The TLS signature also significantly correlated with hallmark scores for hypoxia, IFNα response, IFNγ response, inflammatory response, and angiogenesis in our AML dataset (**Figure 6E**). When looking across the entire TCGA dataset, median TLS score for AML was relatively low (-0.45, rank 24/32), suggesting rarer or subdued TLS signaling programs in AML (**Supplemental Figure 5B**). Diffuse large B-cell lymphoma had the highest median TLS score (0.76), possibly due to aberrant CCR7 expression in malignant B cells^28^.

## Discussion

Uncovering AML-BMM interactions is essential to untangling the complex interplay between leukemia and immunotherapy. Comprehensive characterization of the BMM is necessary to determine cell states and interactions, and requires targeting of tumor, immune, and structural components to uncover deeper spatial characteristics.

AML is a heterogeneous disease, with a diverse range of phenotypic markers that contain a high overlap with normal hematopoietic cells. Currently, no singular marker can confidently phenotype AML blast cells and clinical diagnostic flow cytometry panels for AML exceed 20+ markers to differentiate AML immunophenotypes which carry diagnostic and prognostic implications^29^. Other groups have combined multiple targeted spatial imaging modalities for same slide imaging to improve cell segmentation or augment cell identification and analysis^30–32^. We employed IMC to validate our findings in IF, with the additional benefit of expanding the panel size without subjecting the tissue to additional imaging cycles. By using an adjacent H&E-stained slide as reference, we can align to a minimally treated image that avoids tissue damage or bone detachment from antigen retrieval and multiple staining rounds. Additionally, other structural stains relevant to bone marrow pathology, such as trichrome and silver stains can be performed. Overall, our study finds sequential IF and IMC reliable methods for interrogating the BMM, with good concordance after alignment.

By clustering of the spatial K-function of each cell, we were able to reveal B and T cell aggregates that resembled various maturation states of TLS in solid tumors^33^. Current definitions of TLS are variable, and several delineations of TLS maturity and development have been proposed, with the most immature state being a loose lymphocyte aggregate while the most mature feature an organized cluster of lymphocytes with a germinal center, high endothelial venules, T follicular helper cells and follicular dendritic cells^33^. Lymphocyte aggregates have long been observed in bone marrow, being described on a spectrum from disorganized clusters to structures resembling classical TLS. Their effect on the AML BMM however has not been thoroughly detailed^34–36^. These “TLS-like aggregates” were recently observed in a pediatric AML population and found to be increased in number after immunotherapy^20^.

Interestingly, all of the patients in our study exhibiting TLS-like aggregates (n=3) were positive for TP53 mutation, while their cohort was enriched for KMT2A rearrangement. TP53 is a well-known cancer driver which has been associated with TLS signatures in several cancers and corresponds with poor prognosis in AML^37,38^. TLS are thought to be formed under chronic inflammatory conditions^39^, and similarly, AML progression and pathogenesis has been linked to inflammation^40^. Our previous work implicated HLA-E as an immune evasion reaction of AML cells upregulated in response to inflammatory cytokines such as IFNγ^27^. This finding is corroborated by the strong correlation between HLA-E expression and a validated TLS score in a large, integrated public AML dataset. Our research shows that the transcriptomic signature of these TLS-like aggregates can be observed spatially and corresponds with better survival, similar to that of TLS in solid cancers^23,41,42^. Though in the present study we lack the ability to detect features of mature TLS, our integration of multiple spatial proteomics, spatial transcriptomics, and bulk RNA assays suggests the presence of bona-fide TLS may exist in AML.

Currently, many questions remain unanswered about the state of TLS-like and lymphocyte aggregates in AML. Lymphocyte aggregates in bone marrow are relatively rare in occurrence, estimated to be between 1-13%^43^. Even more rare, Agbay et al. reported less than 0.02% incidence of reactive germinal centers in 205,274 bone marrow specimens, and none in the context of AML^44^. In contrast, TLS are reported to occur in up to 60% of breast cancer cases^45^. Several models of TLS maturation suggest lymphocyte aggregates as the initial steps to TLS formation^33,46^. Though only three aggregates were found in the present study, only one had characteristics resembling a mature TLS. As previous studies have shown that only mature TLS exhibit favorable prognostic value^37,41^, understanding mechanisms of TLS formation and maturation in AML compared to other cancers is crucial. Currently, the value of inducing TLS remains uncertain. The stimulator of interferon genes (STING) agonist ADU S-100 has been reported to promote TLS formation through LTα, IL36β and TNFα in a melanoma mouse model^47^, however, clinical trials of STING agonists have had disappointing results^48^ with companies halting development on STING agonists GSK3745417, MK-1454, and ADU-S100. Additional areas of investigation include prognostic features of TLS which include TLS maturity, TLS density, and intra or peri-tumor location in solid tumors^49^. Though not all these features are directly applicable to liquid malignancies, possible analogs in the AML BMM could include size, proximity, and density of TLS around AML-enriched spatial neighborhoods. Given the potential impact on immune response and scarcity of research on TLS in AML, further study into these TLS-like aggregates, their factors of formation and maturation, their interaction with the BMM, and their effect on immunotherapy is warranted.

In summary, we were able to characterize a TLS-like aggregate in AML patients through an optimized, flexible, and cost-effective pipeline performant enough to spatially interrogate the leukemic bone marrow microenvironment at the single cell level using same slide IF and IMC. We were able to further validate the presence of TLS-like aggregates using established TLS signatures in spatial transcriptomics and public AML datasets and show correlations to prognostic impact. Further research into TLS-like aggregates in the BMM could help unveil spatial factors of durable immune response in AML.

## Materials and Methods

### TMA Preparation

16 bone marrow biopsies from 7 patients were retrieved from Department of Hematopathology archives at the University of Texas MD Anderson Cancer Center and organized into two tissue microarrays (TMAs). Regions of interest (ROIs) for each sample were annotated using the TMA control software to select for focal areas of cellularity (3DHISTECH, Budapest, Hungary). A FFPE TMA block was created by embedding a 1.5 mm diameter circular core from each donor block with 0.5 mm spacing using the TMA Grand Master (3DHISTECH), which was sectioned at 4 µm to create a series of TMA slides. The study protocol was approved by MD Anderson’s institutional Review Board. The study was conducted in accordance with the principles of the Declaration of Helsinki.

### COMET Multiplex IF

FFPE TMA slides underwent dewax and hydration largely as previously described^50^. Slides then underwent heat induced epitope retrieval (HIER) at 107° C for 15 minutes. For testing of the Lunaphore protocol, dewax and antigen retrieval was performed using the PT module (Epredia, Kalamazoo, MI) as previously described^6^. For testing of the TrueBlack, TrueBlack lipofuscin autofluorescence quencher (Biotium Inc, Fremont, CA) was applied according to manufacturer directions. The slide was then loaded into the COMET to fit in the 9×9 mm square imaging window. Staining, imaging, and elution was performed largely as previously described^51^. Primary antibodies and dilutions are found in **Table 2**. Imaged slides were stored in PBS at 4° Celsius until ready to stain for IMC. **Opal multiplex IF**

FFPE TMA slides from the same blocks were baked, dewaxed, and treated with 3% peroxide. Tissue then underwent the 7-color Opal multiplex staining workflow. In short, slides underwent HIER, blocked, and incubated with primary antibodies (**Table 3**). Slides were then incubated with horseradish peroxidase, then Opal fluorophores, stripping the out the antibodies between each marker before counterstaining with 4’,6-diamidino-2-phenylindole (DAPI) and mounting with a coverslip. All steps were performed on the NanoVIP 100 automated stainer (Biogenex, Fremont, CA). Slides were then imaged at 40X using the PhenoImager HT 2.0 (Akoya Biosciences) at 40X. **IMC Staining and Imaging**

Briefly, the TMA slide was treated with blocking buffer and permeabilization reagents, incubated in a cocktail of metal conjugated antibodies, (**Table 4**) and counter stained with iridium intercalator (201192B, Standard BioTools, San Franciso, CA). After drying, ROIs were drawn to select COMET imaged cores, which were imaged at 10X using the Hyperion Xti (Standard BioTools). The resulting MCD files were converted into TIFF by MCD viewer (v1.0.560.6, Standard BioTools).

### Image Analysis

Image files were analyzed using the Visiopharm image analysis software (v2023.09, Visiopharm A/S, Hoersholm, Denmark). COMET IF, IMC, and H&E images were aligned using the TissueAlign function guided by major tissue landmarks and coexpression of markers across assays. Detection of hematopoietic tissue and bone tissue was performed using a random forest classifier on the DAPI stain and the H&E stain, respectively. Areas with necrosis, crushed cells, poor focus, misaligned stitching, or high autofluorescence were removed. COMET image background subtraction was performed by subtracting the autofluorescence captures scaled by the average autofluorescence of previous runs per channel from the marker image. Cell segmentation was achieved by detecting cell nuclei using DAPI or iridium counterstains with a deep learning algorithm pre-trained on a U-Net convolutional neural network. Cells were classified using a supervised gating strategy based on mean intensity thresholds applied to canonical markers while leukemic blasts were annotated using patient specific immunophenotypes obtained by flow cytometry (**Tables 5 and 6**). For validation using the Opal staining workflow, cores with lymphocytes aggregates in the COMET assay were selected for comparison of staining. Cell labels, MFIs, and XY positions were exported.

**Table 5:** COMET IF gating strategy. *AML blasts were gated per patient based on clinical flow cytometry immunophenotype reports

| COMET IF |  |  |  |  |
| --- | --- | --- | --- | --- |
| Cell Type | Antigen | Clone | Clonality | Source |
| B Cell | CD45 (bright) | BL-178-12C7 | Rabbit Monoclonal | Bethyl |
|  | CD20 | L26 | Mouse Monoclonal | Bethyl |
| CD4T Cell | CD45 (bright) | BL-178-12C7 | Rabbit Monoclonal | Bethyl |
|  | CD3 | 3F3A1 | Mouse Monoclonal | ProteinTech |
|  | CD4 | BL-155-1C11 | Rabbit Monoclonal | Bethyl |
| Regulatory T Cell | CD45 (bright) | BL-178-12C7 | Rabbit Monoclonal | Bethyl |
|  | CD3 | 3F3A1 | Mouse Monoclonal | ProteinTech |
|  | CD4 | BL-155-1C11 | Rabbit Monoclonal | Bethyl |
|  | FoxP3 | BLR034F | Rabbit Monoclonal | Bethyl |
| CD8T Cell | CD45 (bright) | BL-178-12C7 | Rabbit Monoclonal | Bethyl |
|  | CD3 | 3F3A1 | Mouse Monoclonal | ProteinTech |
|  | CD8 | C8/144B | Mouse Monoclonal | Bethyl |
| Natural Killer Cell | CD45 (bright) | BL-178-12C7 | Rabbit Monoclonal | Bethyl |
|  | CD56 | BLR171J | Rabbit Monoclonal | Bethyl |
| Monocyte/<br>Macrophage<br>Lineage Cells | CD45(dim) | BL-178-12C7 | Rabbit Monoclonal | Bethyl |
|  | CD14 | BLR187J | Rabbit Monoclonal | Bethyl |
|  | CD68 | KP1 | Mouse Monoclonal | Bethyl |
|  | CD163 | 10D6 | Mouse Monoclonal | Invitrogen |
| AML Blast* | CD45(dim) | BL-178-12C7 | Rabbit Monoclonal | Bethyl |
|  | CD4 | BL-155-1C11 | Rabbit Monoclonal | Bethyl |
|  | CD14 | BLR187J | Rabbit Monoclonal | Bethyl |
|  | CD34 | EP373Y | Rabbit Monoclonal | Abcam |
|  | CD56 | BLR171J | Rabbit Monoclonal | Bethyl |

**Table 6:** IMC gating strategy.

| IMC |  |  |  |  |
| --- | --- | --- | --- | --- |
| Cell Type | Antigen | Clone | Clonality | Source |
| Helper T Cell | CD3 | MRQ-39 | Rabbit Monoclonal | CellMarque |
|  | CD4 | EPR6855 | Rabbit Monoclonal | Fluidigm |
| Cytotoxic T Cell | CD3 | MRQ-39 | Rabbit Monoclonal | CellMarque |
|  | CD8 | D8A8Y | Rabbit Monoclonal | Fluidigm |
| Monocyte/<br>Macrophage<br>Lineage Cells | CD14 | EPR3653 | Rabbit Monoclonal | Fluidigm |
|  | CD68 | KP1 | Mouse Monoclonal | Fluidigm |
|  | CD163 | BLR087G | Mouse Monoclonal | Bethyl |
| AML Blast | CD34 | QBEND/10 | Mouse Monoclonal | Invitrogen |

### Visium

Briefly, FFPE bone marrow samples were processed using the standard CytAssist protocol from 10X Genomics. SpaceRanger (v2.0) was used to align sequences to the reference genome and to map spots to the tissue. Spots were filtered (See Supplemental Methods), normalized using SCTransform, and variable genes identified using the FindVariableFeatures function. Cell deconvolution used the Seurat transfer learning pipeline leveraging in-house single-cell RNA-seq data. TLS signatures were scored using the AUCell R package^52^.

### Bulk data

RNA-seq data and corresponding clinical data from TCGA, BeatAML and MDACC were downloaded and integrated into one dataset. Batch effects were corrected using ComBat-seq^53^. TLS signatures obtained from the literature were used to score the data using the R package GSVA^54^.

### Statistical Analysis

All statistical analyses were performed in the R statistical software (v4.3.0) using the rstatix R package^55^. Marker expression was arcsinh transformed and analyzed using the imcRtools package^56^. Spatial regions were generated with the lisaClust package^57^. Cell distances were calculated using the minDistToCells() function. Two-sided Wilcoxon rank sum test was used for comparisons between two independent groups while the paired Wilcoxon was used for paired data. Survival curves were created using the Kaplan-Meier method and assessed using the log-rank test. Pearson correlation was used for all correlative tests. All statistical tests used an alpha of 0.05.

### Data visualization

Pipeline workflow was created in BioRender (BioRender, Toronto, Canada). Figures were constructed using the ComplexHeatmap^58^, pheatmap^59^, ggplot2^60^, ggpubr^61^, and dittoSeq^62^ R packages and composed in Illustrator (v28.1, Adobe; San Jose, CA). All boxplots use median for the center line Micrographs were created by cropping original image files then layering and applying pseudocolor in Photoshop (v25.5.1, Adobe) through linear mapping. Whole image brightness adjustments (levels) were made for visibility.

Extended details of the methods and materials used can be found in **Supplemental Methods**.

## Data Availability

The datasets generated during the current study will be available upon publication. Additionally, COMET images will be available through an interactive viewer^63^. The following public datasets were used in this publication and can be accessed at their respective locations: BEAT-AML (https://www.nature.com/articles/s41586-018-0623-z#Sec38), TCGA (https://gdc.cancer.gov/about-data/publications/pancanatlas), MD Anderson (GEO accession CSE165656).

## CODE AVAILABILITY

The underlying code for this study is available in GitHub and can be accessed via this link http://github.com/abbaslab/.

## Acknowledgements

We would like to thank Thomas Huynh and Arizona Nguyen at the Department of Veterinary Services at MD Anderson for their work and consultation on TMA design. This study received no funding.

## Author Contributions Statement

Contribution: H.A.A. devised and directed the project; C.P.L. and I.V. co-analyzed the data. E.D. performed the spatial transcriptomics analysis. C.P.L. wrote the manuscript with contributions from H.A.A., I.V., S.F.B., P.K.R. and J.K.B.; A.V.B and J.A.G. assisted in the acquisition and optimization of IMC and IF imaging; J.L.R. performed IMC antibody conjugation; C.P.L performed the image analysis and quality check with assistance from C.D.P. and I.V.; F.Z.J. and A.E.Q. assisted in panel design and sample collection with C.B.R. and M.G; P.S and S.B. provided assistance in preparation and analysis of the Visium assay. All authors provided feedback and contributed to the final manuscript.

