## Supplemental Materials for "Multimodal Spatial Proteomic Profiling in Acute Myeloid Leukemia"

### Supplemental Methods

#### TMA Preparation

16 bone marrow biopsies from 7 patients were retrieved from Department of Hematopathology archives at the University of Texas MD Anderson Cancer Center and organized into two tissue microarrays (TMAs). Regions of interest (ROIs) for each sample were annotated using the TMA control software to select for focal areas of cellularity (3DHISTECH, Budapest, Hungary). A FFPE TMA block was created by embedding a 1.5 mm diameter circular core from each donor block with 0.5 mm spacing using the TMA Grand Master (3DHISTECH), which was sectioned at 4 µm to create a series of TMA slides. The study protocol was approved by MD Anderson's institutional Review Board. The study was conducted in accordance with the principles of the Declaration of Helsinki.

#### Sample preparation

FFPE (formalin-fixed, paraffin embedded) TMA slides were baked for 60 minutes at 60° Celsius, dewaxed 3 times using xylene (214736-1L, Sigma-Aldrich, St Louis, MO) for 5 minutes each, and rehydrated in a graded ethanol series of 100% ethanol for 6 minutes, 95% ethanol for 6 minutes, 70% ethanol for 6 minutes, 50% ethanol for 6 minutes, 30% ethanol for 6 minutes, then PBS (AM9625, Invitrogen, Waltham, MA) for 2 minutes. Slides were then treated with 3% hydrogen peroxide (216763-100ML, Sigma-Aldrich) for 10 minutes before being washed with PBS. Antigen retrieval was performed with an EZ-Retriever system V.3 (BioGenex, Fremont, CA) at 107° C for 15 minutes in an EDTA (Ethylenediaminetetraacetic acid) based buffer (EZ-AR 2 Elegance, HX032YCX-GP, Biogenex).

For testing of the TrueBlack protocol, sample preparation proceeded as previously mentioned. TrueBlack lipofuscin autofluorescence quencher 20X (#23007, Biotium) was then warmed in a water bath to 70° C for 15 minutes. The solution was then diluted to 1X in 70% ethanol before being incubated on the TMA slide for 1 minute then washing the slide three times in PBS.

For testing of the standard Lunaphore protocol, deparaffinization and antigen retrieval was performed using the PT module (Epredia, Kalamazoo, MI) as previously described[1] using Dewax and HIER buffer H (TA-999-DHBH, Epredia) for 60 minutes at 102° C and stored in PBS (AM9625, Invitrogen) until use.

Comparison of the standard Lunaphore protocol and our optimized TrueBlack protocol revealed higher autofluorescence intensities with comparable marker intensities for the standard Lunaphore protocol (**Supplemental Figure 1C-F**). As such we used our protocol with TrueBlack autofluorescence quenching for our analysis.

**COMET Multiplex IF**

The Lunaphore COMET sequential IF platform was used to profile the leukemic and immune bone marrow microenvironment. Primary antibodies were diluted in multistaining buffer (BU06, Lunaphore Technologies). Secondary anti-rabbit/anti-mouse Alexa Fluor 555 and 647 (A21430, A48289, Invitrogen) were also diluted in multistaining buffer at 1:100 and 1:200 dilutions respectively in conjunction with 4',6-diamidino-2-phenylindole (DAPI) (62248, Thermo Fisher Scientific, Waltham, MA) at a 1:2000 dilution for the nuclear counterstain. List of primary antibodies with corresponding incubation times and dilutions are found in **Supplementary Table 2**. Each slide was then loaded into the COMET to fit in a 9×9 mm square imaging window. Staining, imaging, and elution was performed cyclically by the COMET using the "Characterization 2" protocol to capture an unstained autofluorescence image for each channel in every cycle. For each cycle the following settings were used: Incubation time was 4 minutes for primary antibodies and 2 minutes for secondary antibodies. Exposure times used for each channel were: DAPI 80 *ms*, TRITC 400 *ms*, Cy5 200 *ms*. The elution step lasted 4 min for each cycle and was performed with Elution Buffer (BU07-L, Lunaphore Technologies) at 37°C. Quenching was performed with Quenching Buffer (BU08-L, Lunaphore Technologies). Imaging step was performed at 20X with Imaging Buffer (BU09, Lunaphore Technologies). Stitching, alignment, and flat field correction was automatically performed by the COMET system and an OME.TIFF file was produced. Slides were then stored in PBS at 4° Celsius until they were ready to stain for IMC.

**Opal multiplex IF**

FFPE TMA slides from the same blocks were baked at 60° Celsius overnight. Slides were then placed on the NanoVIP 100 automated stainer (Biogenex) and a protocol created to execute the following steps. Slides were first washed with X-Dewax solution (HX016-XEK, Biogenex) 8 times in 2 minute intervals before being washed twice with PBS (P2100-100, GenDEPOT, Katy, TX). Slides were then incubated in 3% peroxide (Thermo Fisher Scientific, BP2633-500) for 15 minutes. HIER was performed with EZAR 1 (HK546-XAK, Biogenex) for 15 minutes at 95° Celsius the washed twice with PBS. Blocking was performed with 2.5% goat serum (S-1012, Vector Laboratories, Newark, CA) for 20 minutes. Antibodies diluted in antibody diluent (S0809, Agilent, Santa Clara, CA) were incubated for 1 hour before washing 3 times with Tween 20 (P7949, Sigma-Aldrich) then incubated in either goat anti-rabbit or goat anti-mouse horseradish peroxidase polymer (MP-7451, MP7452, Vector Laboratories) and finally washed 3 times with Tween 20. Slides were incubated in Opal fluorophores diluted in amplification diluent (FP1609, Akoya Biosciences, Marlborough, MA) for 10 minutes before washing 3 times with Tween 20. Antibodies were stripped using EZ-AR1 incubation for 15 minutes at 95° Celsius and washed twice with PBS. This was repeated for each antibody before incubating with in a DAPI counterstain for 2 minutes and washing twice with PBS. Mounting medium (H-1700, Vector Laboratories) was applied and coverslipped. Slides were then imaged using the PhenoImager HT 2.0 (Akoya Biosciences) at 40X.

**IMC staining and imaging**

The TMA slide was incubated in a cocktail of metal conjugated antibodies (**Supplementary Table 2**) diluted in blocking buffer consisting of 2% BSA (1000161,

Jackson Immuno Research Labs, West Grove, PA) and 0.5% NHS (S-2000, Vector Laboratories, Newark, CA) overnight at 4° Celsius. The slides were then treated with 0.2% TritonX-100 (x100, Sigma-Aldrich) in DPBS (P2100-100, GenDEPOT) twice for 10 minutes, then TBS (T8057-100, GenDEPOT) twice for 10 minutes. Intercalator (201192B, Standard BioTools, San Franciso, CA) stain was performed with a 1:400 dilution in DPBS for 30 minutes followed by a wash in ddH2O (400000, Cayman Chemical) for 5 minutes before air drying. After drying, ROIs were drawn to select 2 Lunaphore imaged cores on TMA7, which were imaged at 10X using the Hyperion Xti (Standard BioTools). The resulting MCD files were converted into TIFF by the MCD viewer software (v1.0.560.6, Standard BioTools).

**Imaging Mass Cytometry QA**

IMC quality and concordance with IF was validated by assessing the quality of nuclear stain on IMC at different time points (47 and 71 days) post IF imaging (**Supplemental Figure 3A**). Cell segmentation from both modalities was overlaid and compared. The lower quality of cell segmentation on IMC impelled to use the DAPI nuclear stain for aligned images moving forward (Supplemental Figure 3B).

**Background Subtraction**

Autofluorescence is a major obstacle in bone marrow tissue as high levels of autofluorescence in the 450-550 nm range[2] emerge due to lipofuscin, red blood cells, and extracellular matrix components such as collagen and elastin[3–5]. To abrogate this, we captured the autofluorescence intensities in conjunction with the stained mean fluorescence intensities (MFI) of each marker for each cell. The Lunaphore background subtraction algorithm subtracts out the initial first cycle autofluorescence capture from every stained image. Since autofluorescence decreases each cycle with repeated application of the quenching buffer, this results in over-subtraction for later cycles (**Supplemental Figure 1A**). We adjusted for this effect by adjusting the magnitude of the subtraction by the average TRITC per cycle to obtain a scaled subtraction effect that more accurately removed background noise (Supplemental Figure 1B). TIFF files were imported into the Visiopharm image analysis software (v2023.09, Visiopharm A/S, Hoersholm, Denmark). Each marker image was subtracted by the autofluorescence feature, which was reduced by a fixed multiplier depending on the cycle. This multiplier was derived from historical runs where autofluorescence MFI was measured per cycle. These post-subtraction features were then used as expression for downstream analysis.

**Image Analysis**

COMET IF, IMC, and H&E images were then aligned using the TissueAlign function guided by major tissue landmarks and coexpression of markers targeting the same antigen across assays. Detection of hematopoietic tissue and bone tissue was performed using a random forest classifier on the DAPI stain and the H&E stain, respectively. Areas with necrosis, crushed cells, poor focus, misaligned stitching, or high autofluorescence were removed. Cell segmentation was achieved by detecting cell nuclei using DAPI or iridium counterstains with a deep learning algorithm pre-trained on a U-Net convolutional neural network. Subsequently, cell boundaries were identified by extending outward from the nuclear 'seeds' based on predictions of cytoplasmic areas. Cells were classified using a supervised gating strategy based on mean intensity thresholds applied to canonical markers while leukemic blasts were annotated using patient specific immunophenotypes obtained by flow cytometry. For validation using the

Opal staining workflow, cores with lymphocytes aggregates in the COMET assay were selected for comparison of staining. Cell labels, MFIs, and XY positions were exported.

**Bulk Transcriptomics**

RNA-seq data and corresponding clinical data from BEAT-AML (https://www.nature.com/articles/s41586-018-0623-z#Sec38), TCGA (https://gdc.cancer.gov/about-data/ publications/pancanatlas), MD Anderson (GEO accession CSE165656) were downloaded and integrated into one dataset. Batch effects were corrected using ComBat-seq[6]. Analysis was limited to bone marrow samples in adult patients with newly diagnosed AML. TLS signatures were obtained from the literature[7–11] (See Table 4) and gene set variation analysis was performed using the R package GSVA[12]. Survival analysis was performed using the survminer package[13]. After dichotomizing signatures scores and HLA-E expression by the median, survival curves were created using the Kaplan-Meier method and comparison between high and low groups was assessed with the log-rank test. Patients were right censored at 5 years. For HLA-E correlation to TLS score, 4 extreme outliers of HLA-E expression (around 33000 fold the median value) were removed from the MDACC dataset after outlier detection using the iterative Grubbs test.

**Statistical Analysis**

All statistical analyses were performed using either the base R statistical software (v4.3.0) or rstatix R package[14]. Exported data from Visiopharm packaged into the SpatialExperiment container using the imcRtools pipeline[15]. Marker expression was transformed by the arcsinh function. Principle component analysis was done using the runPCA function from the scater package[16]. Integration and batch correction was done using Harmony[17], and the Harmony embeddings were placed into low-dimensional space using runUMAP and runTSNE functions. Spatial regions were analyzed with the lisaClust package[18] by performing K-means clustering on the curves generated from the K function at radii of 10, 20, 50, 80, and 120 μm and k = 5 centroids. Cell distances to were calculated using the minDistToCells() function of the imcRtools package. The two-sided Wilcoxon rank sum test was used for all comparisons between two independent groups. For paired comparisons the two-sided paired Wilcoxon rank sum test was used. Pearson correlation was used for all correlative tests. All statistical tests were calculated using a p value of $p < 0.05$ for significance.

**Spatial Transcriptomics**

Formalin-fixed paraffin-embedded (FFPE) tissue from bone marrow biopsies was used. RNA integrity was assessed by calculating DV200 values using an Agilent Bioanalyzer. The samples were processed using the standard Visium CytAssist Spatial Gene Expression protocol from 10X Genomics. The Seurat pipeline was applied for downstream analysis[19]. The median value of the mitochondrial read percentage was determined and spots filtered by setting minimum and maximum thresholds at ±3 times the median absolute deviation (MAD): $median(|x - median(x)|)$ where $x$ is the spatial transcriptomics (ST) feature. The same calculation was applied to the log10 values of the number of genes captured, while for the number of oligonucleotides, the algorithm was used only for setting maximum thresholds. Spots that fell within these thresholds were retained for subsetting the ST data. Filtered spots were then normalized using SCTransform. The top 5000 variable genes were identified using the FindVariableFeatures function with variance stabilizing transformation. Principal component analysis was performed using the RunPCA function, and Uniform Manifold Approximation and Projection layouts and nearest-neighbor graphs were identified using the top 25 principal components. Our spatial transcriptomics data were deconvolved using a transfer learning pipeline in Seurat, leveraging in-house generated single-cell RNA-seq data from healthy (n=9) and AML-affected (n=7) bone marrow suspensions. TLS signatures were scored on a spot-by-spot basis using the AUCell algorithm[20]. Hotspots with high TLS signal were selected using 10X Genomics' Loupe Browser (version 7.0). Pearson correlation of the deconvolution results for TLS signatures, B cells, and CD4 Effector cells within the selected region was calculated.

**Data visualization**

Pipeline workflow was created in BioRender (BioRender, Toronto, Canada) and edited in Illustrator (v28.1, Adobe; San Jose, CA, USA). Heat maps were constructed using the ComplexHeatmap[21] and pheatmap[22] R packages. Plots were constructed using the ggplot2[23], ggpubr[24], and dittoSeq[25] R packages and composited in Illustrator. Micrographs were created by cropping the original image files then layering and applying pseudocolor in Photoshop (v25.5.1, Adobe). Whole image brightness adjustments (levels) were edited for visibility.

### Supplemental Figures

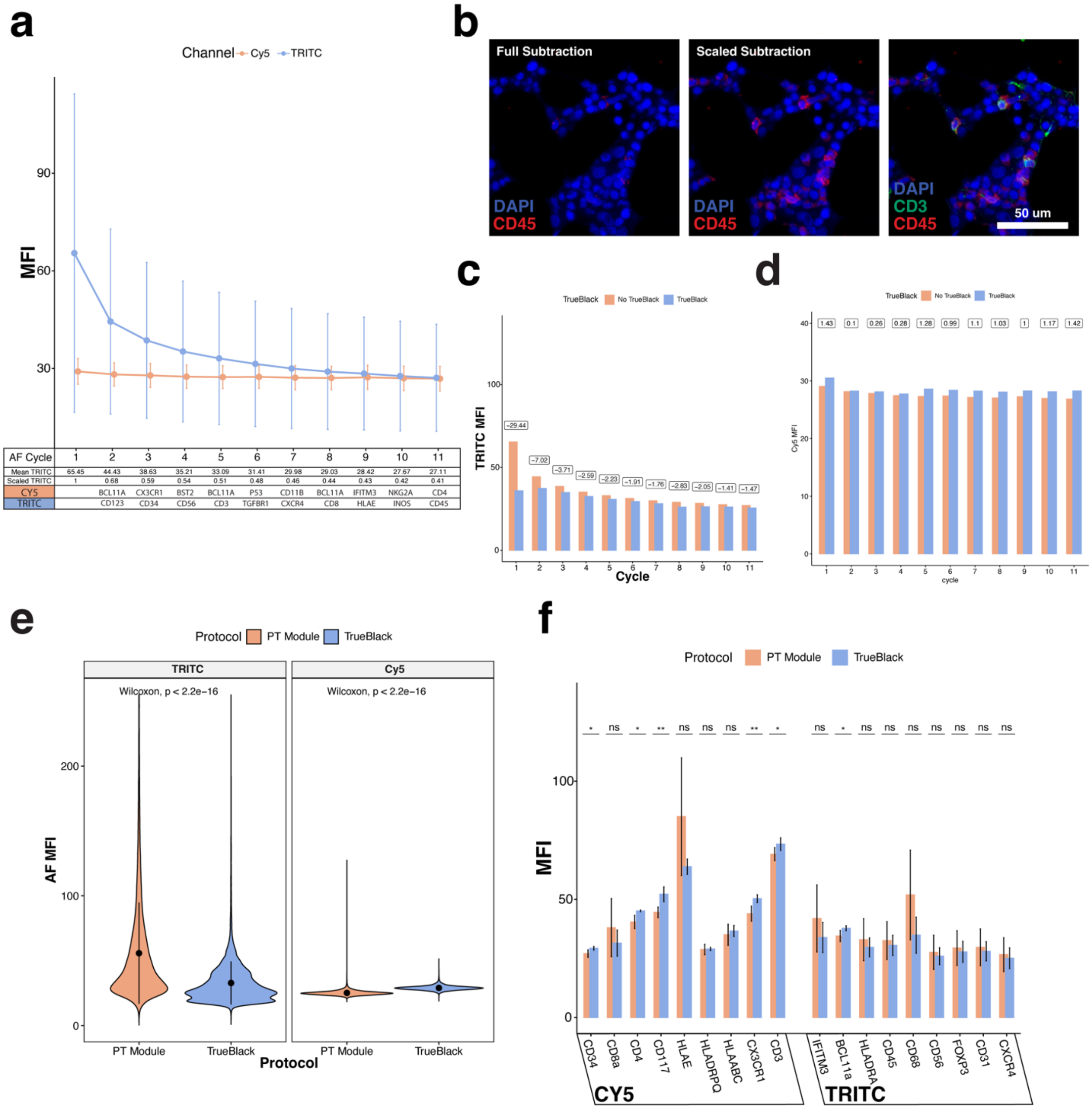

**Supplemental Figure 1: Autofluorescence in the COMET sequential IF system can be abrogated by a scaled subtraction algorithm. (**A) Line graph shows early rapid decrease in 555 autofluorescence in early imaging cycles. (B) Scaling autofluorescence subtraction by cycle recovers quenched signal in CD45+ cells. (C) Application of TrueBlack reduces average 555 autoflourescence by 45% in the first imaging cycle before aligning with untreated tissue. Boxes show average difference in MFI per cycle. (D) Application of TrueBlack has little effect on 555 markers and moderate increase in autofluorescence for 647 markers. Boxes show average difference in MFI per cycle. (E) Trueblack application results in reduced autofluorescence in the 555 channel and a smaller increase in autofluorescence in 647 channel (Wilcoxon test, two-sided, p <2.2e-16). (F) Trueblack application had insignificant changes in marker intensity for 8/9 555 markers and 4/9 647 markers. Lines denote standard deviation (Wilcoxon Test, two-sided, *=p<0.04, **=p<0.01, ns=not significant).

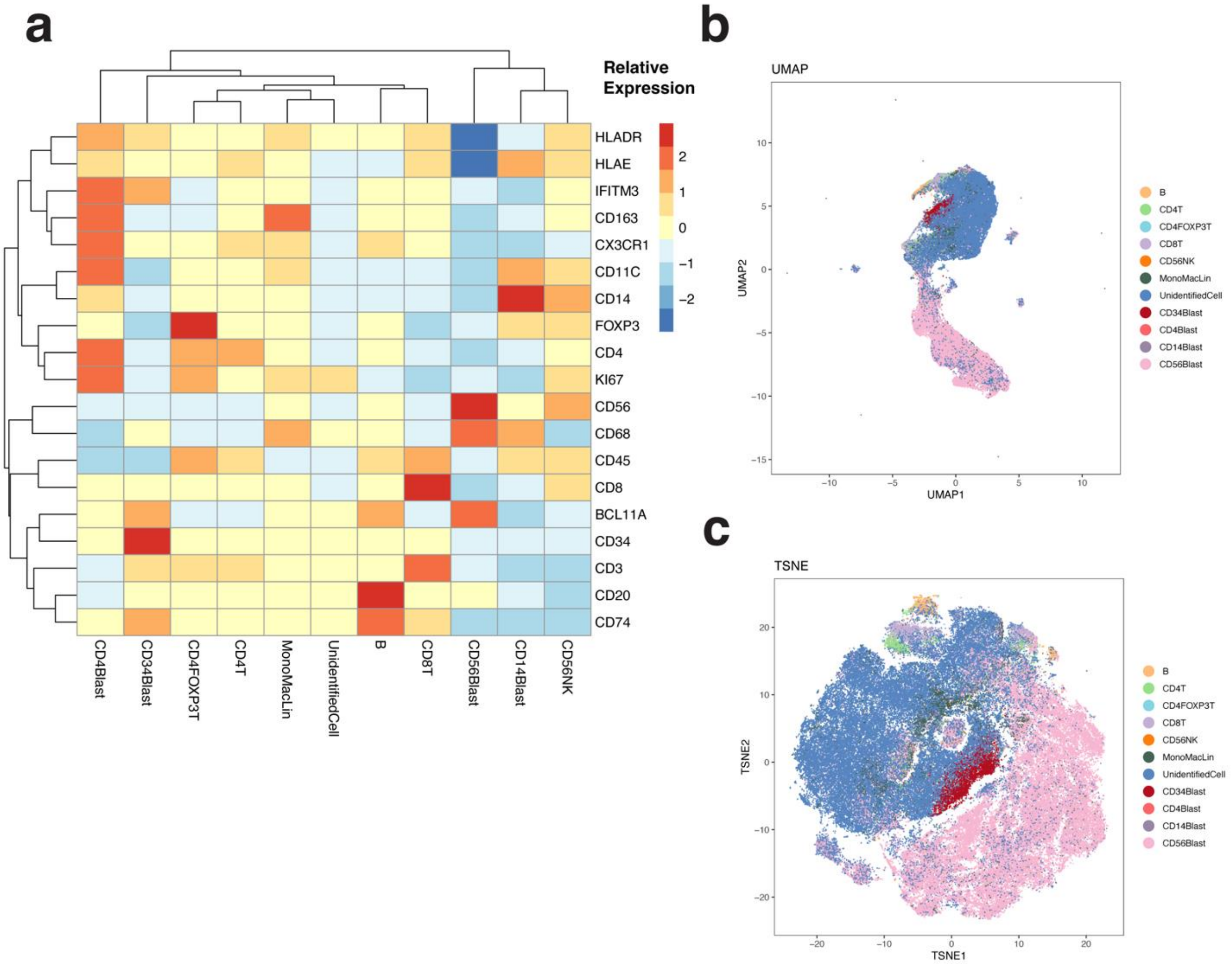

**Supplemental Figure 2: Cell expression characteristics.** (A) Relative expression of cell types on the COMET IF assay. (B) Cellular protein expression projected on to UMAP dimensional space (C) Cellular protein expression projected on to TSNE dimensional space.

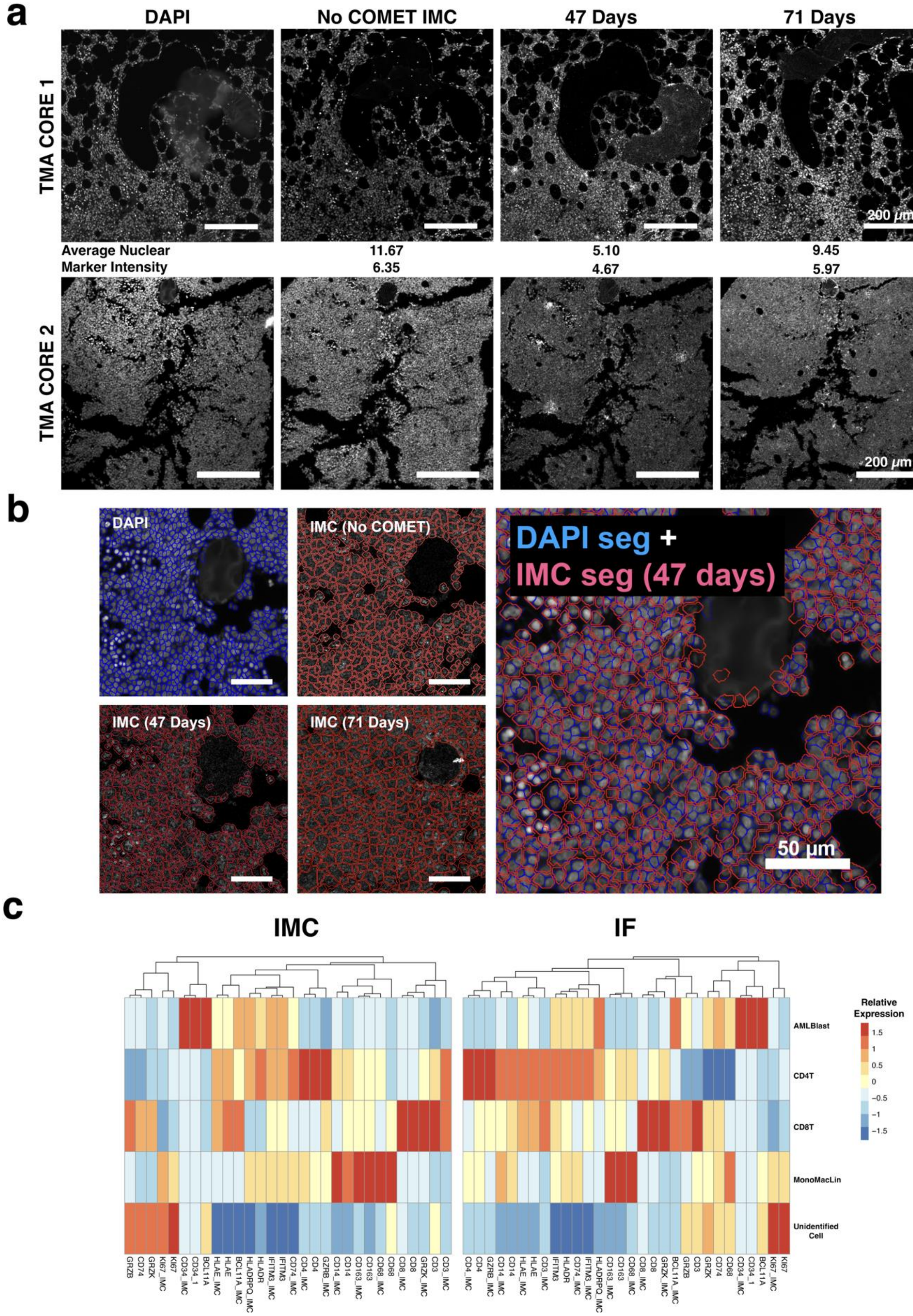
a
DAPI
No COMET IMC
47 Days
71 Days
TMA CORE 1
200 μm
Average Nuclear
Marker Intensity
11.67
6.35
5.10
4.67
9.45
5.97
TMA CORE 2
200 μm
b
DAPI
IMC (No COMET)
IMC (47 Days)
IMC (71 Days)
DAPI seg +
IMC seg (47 days)
50 μm
c
IMC
IF
Relative Expression
AMLBlast
CD4T
CD8T
MonoMacLin
Unidentified Cell

**Supplemental Figure 3: Quality assessment of COMET to IMC staining shows comparable marker staining but degraded nuclear staining.** (A) Nuclear staining of IMC post-COMET staining at 47 days and 71 days compared to IF DAPI stain and IMC staining without COMET. (B) Comparison of cell segmentation on the DAPI stain vs IMC stains post-COMET staining. Segmentation masks were exported from Visiopharm and layered in photoshop. (C) Relative protein expression of cell type labels generated from IMC data vs IF data.

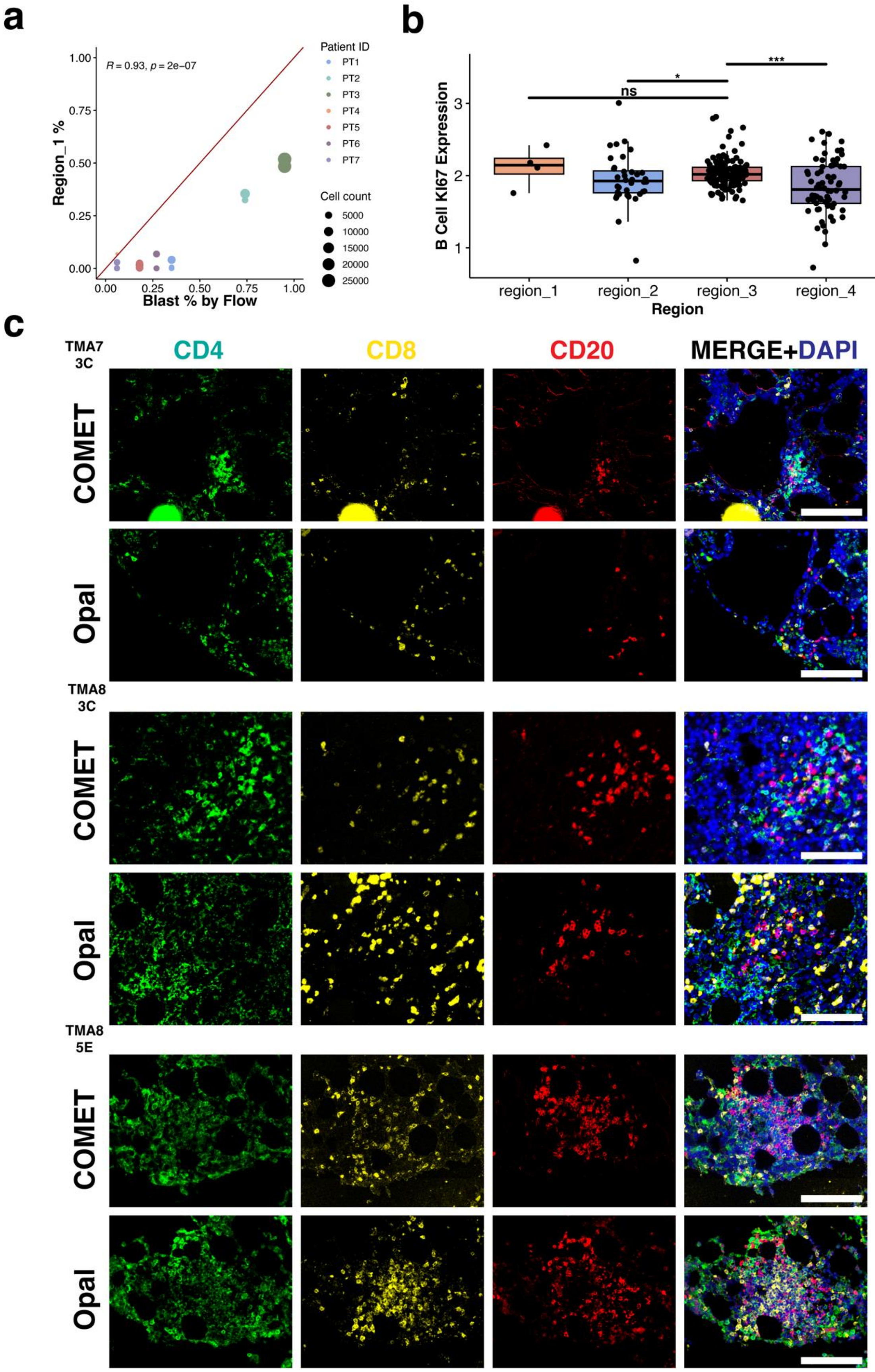
a
R = 0.93, p = 2e−07
Region_1 %
Blast % by Flow
1.00
0.75
0.50
0.25
0.00
0.00
0.25
0.50
0.75
1.00
Patient ID
PT1
PT2
PT3
PT4
PT5
PT6
PT7
Cell count
5000
10000
15000
20000
25000
b
B Cell KI67 Expression
Region
region_1
region_2
region_3
region_4
ns
*
***
c
TMA7 3C
TMA8 3C
TMA8 5E
CD4
CD8
CD20
MERGE+DAPI
COMET
Opal

**Supplemental Figure 4: Lymphocyte aggregates have characteristics of TLS and are validated though the Opal multiplex IF workflow.** (A) Percentage size of AML blast enriched region_1 correlates strongly with blast percentage by clinical flow cytometry. The red line denotes perfect correlation (Pearson, $R = 0.93$, $p=2e-07$). (B) In patients with a lymphocyte aggregate, B cells in TLS-like region_3 have significantly higher Ki67 expression than b cells in other regions. (Wilcoxon test, two-sided, *=$p<0.05$, ***=$p<0.001$, ns = not significant). (C) Micrographs showing each lymphocyte aggregate from COMET staining and Opal staining. Scale bar = 100 μm

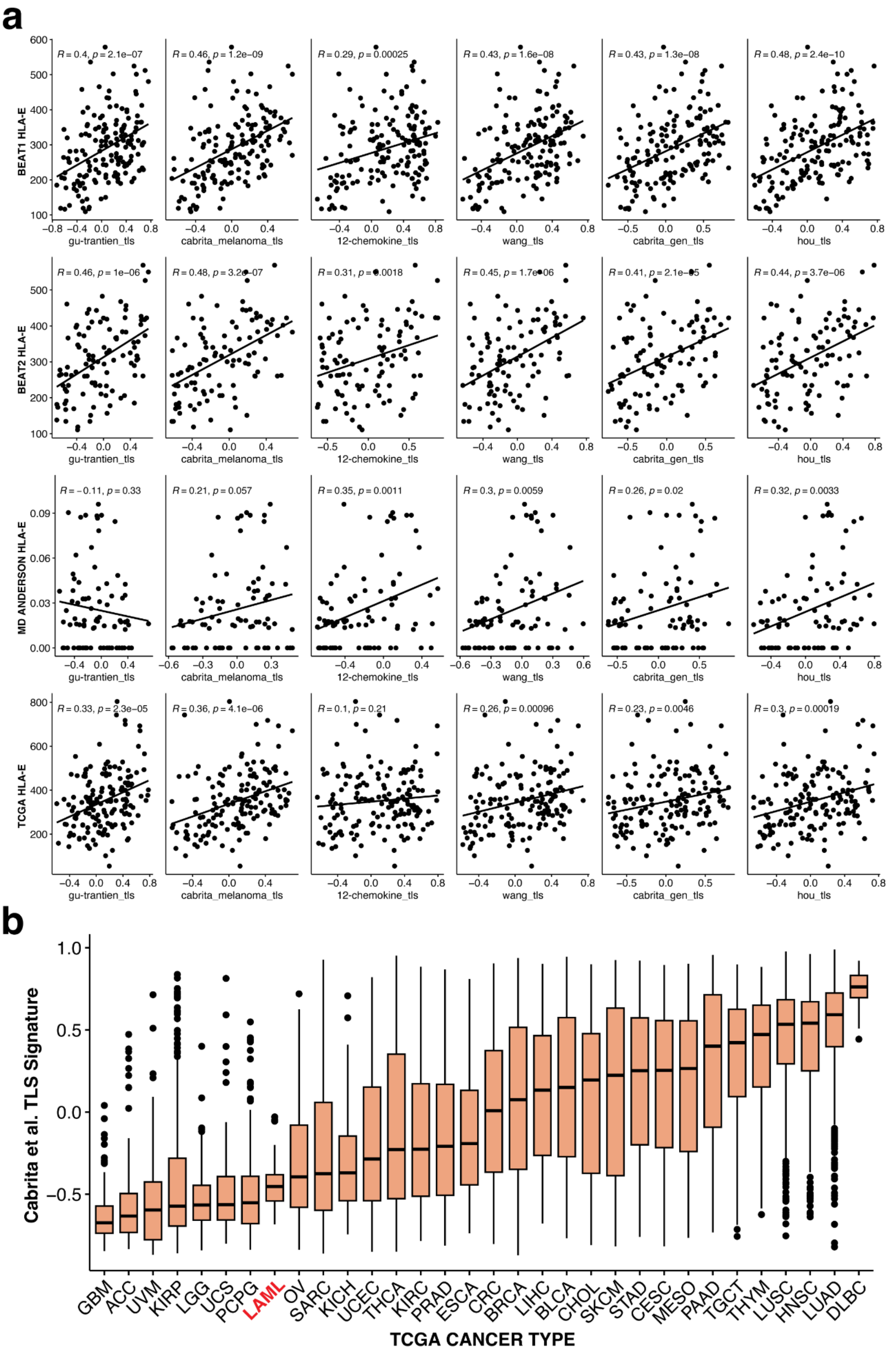

a
BEAT1 HLA-E
BEAT2 HLA-E
MD ANDERSON HLA-E
TCGA HLA-E
gu-trantien_tls
cabrita_melanoma_tls
12-chemokine_tls
wang_tls
cabrita_gen_tls
hou_tls
R = 0.4, p = 2.1e−07
R = 0.46, p = 1.2e−09
R = 0.29, p = 0.00025
R = 0.43, p = 1.6e−08
R = 0.43, p = 1.3e−08
R = 0.48, p = 2.4e−10
R = 0.46, p = 1e−06
R = 0.48, p = 3.2e−07
R = 0.31, p = 0.0018
R = 0.45, p = 1.7e−06
R = 0.41, p = 2.1e−05
R = 0.44, p = 3.7e−06
R = −0.11, p = 0.33
R = 0.21, p = 0.057
R = 0.35, p = 0.0011
R = 0.3, p = 0.0059
R = 0.26, p = 0.02
R = 0.32, p = 0.0033
R = 0.33, p = 2.3e−05
R = 0.36, p = 4.1e−06
R = 0.1, p = 0.21
R = 0.26, p = 0.00096
R = 0.23, p = 0.0046
R = 0.3, p = 0.00019
b
Cabrita et al. TLS Signature
TCGA CANCER TYPE
GBM ACC UVM KIRP LGG UCS PCPG LAML OV SARC KICH UCEC THCA KIRC PRAD ESCA CRC BRCA LIHC BLCA CHOL SKCM STAD CESC MESO PAAD TGCT THYM LUSC HNSC LUAD DLBC

**Supplemental Figure 5: HLA-E correlates with TLS signatures in bulk RNA.** (A) HLA-E expression correlates with TLS signatures from the literature in large public AML datasets (Pearson correlation test). (B) Cabrita et al. TLS signature score in all cancers in the TCGA dataset. AML dataset is highlighted in red.

### Supplemental Table 1: TLS signatures applied to bulk RNA dataset

| TLS Signatures | | |
|---|---|---|
| **Author** | **Genes** | **Tissue** |
| Cabrita et al., 2020 | CCL19, CCL21, CXCL13, CCR7, CXCR5, SELL, LAMP3 | Nonspecific |
| Cabrita et al., 2020 | CD79B, CD1D, CCR6, LAT, SKAP1, CETP, EIF1AY, RBP5, PTGDS | Melanoma |
| Coppola et al. 2011 | CCL2, CCL3, CCL4, CCL5, CCL8, CCL18, CCL19, CCL21, CXCL9, CCL10, CCL11, CCL13 | Colorectal Cancer |
| Gu-Trantien et al., 2013 | CD200, CXCL13, FLJ37440, ICOS, SGPP2, SH2D1A, VSTM3, PDCD1 | Breast Cancer |
| Hou et al., 2022 | CETP, CCR7, SELL, LAMP3, CCL19, CXCL9, CXCL10, CXCL11, CXCL13 | Ovarian Cancer |
| Wang et al, 2024 | CCL2, CCL3, CCL4, CCL5, CCL8, CCL18, CCL19, CCL21, CXCL9, CXCL10, CXCL11, CXCL13, CD79B, CD1D, CCR6, LAT, SKAP1, CETP, EIF1AY, RBP5, PTGDS | Colorectal Cancer |
